# Calm on the surface, dynamic on the inside. Molecular homeostasis in response to regulatory and metabolic perturbation of *Anabaena* sp. PCC 7120 nitrogen metabolism

**DOI:** 10.1101/2020.07.17.206227

**Authors:** Giorgio Perin, Tyler Fletcher, Virag Sagi-Kiss, David C. A. Gaboriau, Mathew R. Carey, Jacob G. Bundy, Patrik R. Jones

## Abstract

Nitrogen is a key macro-nutrient required for the metabolism and growth of biological systems. Although multiple nitrogen sources can serve this purpose, they are all converted into ammonium/ammonia as a first step of assimilation. It is thus reasonable to expect that molecular parts involved in the transport of ammonium/ammonia across biological membranes (i.e. catalysed by AMT transporters) connect with the regulation of both nitrogen and central carbon metabolism. In order to test this hypothesis, we applied both (1) genetic (i.e. Δ*amt* mutation) and (2) environmental treatments to a target biological system, the cyanobacterium Anabaena sp. PCC 7120. Cyanobacteria have a key role in the global nitrogen cycle and thus represent a useful model system. The aim was to both (1) perturb sensing and low-affinity uptake of ammonium/ammonia and (2) induce multiple inner N states, followed by targeted quantification of key proteins, metabolites and enzyme activities, with experiments intentionally designed over a longer time-scale than the available studies in literature. We observed that the absence of AMT transporters triggered a substantial response at a whole-system level, affecting enzyme activities and the quantity of both proteins and metabolites, spanning both N and C metabolism. Moreover, the absence of AMT transporters left a molecular fingerprint indicating N-deficiency even under N replete conditions (i.e. greater GS activity, lower 2-OG content and faster nitrogenase activation upon N deprivation). Contrasting with all of the above dynamic adaptations was the striking near-complete lack of any externally measurable phenotype (i.e. growth, photosynthesis, pigments, metabolites). We thus conclude that this species evolved a highly robust and adaptable molecular network to maintain homeostasis, resulting in substantial internal but minimal external perturbations. The analytical data highlights several internal adaptations, including increased N assimilation (i.e. greater GS activity) and nitrogenase activity (i.e. faster activation upon N deprivation) together with altered amino acids metabolism, as indicated by changes in Gln, Glu and 2-OG, indicating an altered C/N balance. The analyses provides evidence for an active role of AMT transporters in the regulatory/signalling network of N metabolism in this biological system, and the existence of a novel fourth IF7A-independent regulatory mechanism controlling GS activity.

## Introduction

Cyanobacteria are a group of morphologically diverse oxygenic photosynthetic bacteria (Singh & Montgomery, 2011) almost ubiquitous to every habitat on Earth, from hot springs to Antarctic rocks (Percival & Williams, 2013). They are often found as integral members of complex ecosystems representing all three domains of life (Adams & Duggan, 2008; Adams *et al*, 2013) where they contribute to whole ecosystem functionality by photosynthesis-driven assimilation of nutrients. One of the key nutrients they assimilate and provide to the (local) ecosystem is nitrogen (N), an essential building block for amino and nucleic acid biosynthesis. Cyanobacteria have a variety of complementary N assimilatory pathways, including ammonium [NH_4_^+^, (Montesinos *et al*, 1998)], nitrate [NO_3_^-^, (Omata *et al*, 1993)] nitrite [NO_2_^-^, (Bird & Wyman, 2003)] and urea (Valladares *et al*, 2002), and have a key role in the global nitrogen cycle (Flores & Herrero, 2005). Some genera even use amino acids [e.g. arginine and glutamine (Montesinos *et al*, 1997)] or directly fix atmospheric nitrogen [biological nitrogen fixation (BNF), (Herrero *et al*, 2001)], globally contributing 200 million tonnes of fixed N per year (Rascio & La Rocca, 2013). Cyanobacteria are also involved in symbiotic associations, with reduced carbon delivered to cyanobacteria in order to sustain BNF (Backer *et al*, 2018). An example of such symbiotic associations is the aquatic fern *Azolla caroliniana*, which receives fixed N from a filamentous cyanobacterium (*Anabaena azollae*) hosted in the ovoid cavities of the plant’s leaves (Lechno-Yossef & Nierzwicki-Bauer, 2005).

In its free-living form, this cyanobacterium makes a significant contribution to the carbon and nitrogen economy of multiple ecosystems (Kellar & Goldman, 1979). *Anabaena sp. PCC 7120* (henceforth 7120) is an isolated strain showing high genome sequence similarity with *Anabaena azollae* and is commonly used as a model organism to investigate cyanobacterial N-fixation (Herrero *et al*, 2016). As nitrogenases are oxygen-sensitive, photosynthesis-driven BNF calls for spatial and/or temporal separation between the metabolic pathway fuelling energy/carbon inputs (i.e photosynthesis) and N -fixation (Fig. 1). Under diazotrophic conditions, 7120 differentiates 5-10% of its cells into specialised N -fixing heterocysts, following a highly regulated developmental pattern [i.e. a single heterocyst every 10-20 cells (Kumar *et al*, 2010)]. Heterocysts undergo a deep metabolic and structural remodelling to enable efficient N fixation (Golden & Yoon, 1998). The oxygen-evolving photosystem II (PSII) is dismantled, carbon fixation is avoided, photorespiratory activity is increased during differentiation (Valladares *et al*, 2007) and cells are surrounded by a thicker cell envelope [through the deposition of two additional envelope layers, i.e. an inner glycolipids and an outer polysaccharides layer (Nicolaisen *et al*, 2009)] than the vegetative cells, providing the required microoxic environment for N fixation activity (Kumar *et al*, 2010)(Fig. 1).

**Figure 1.**
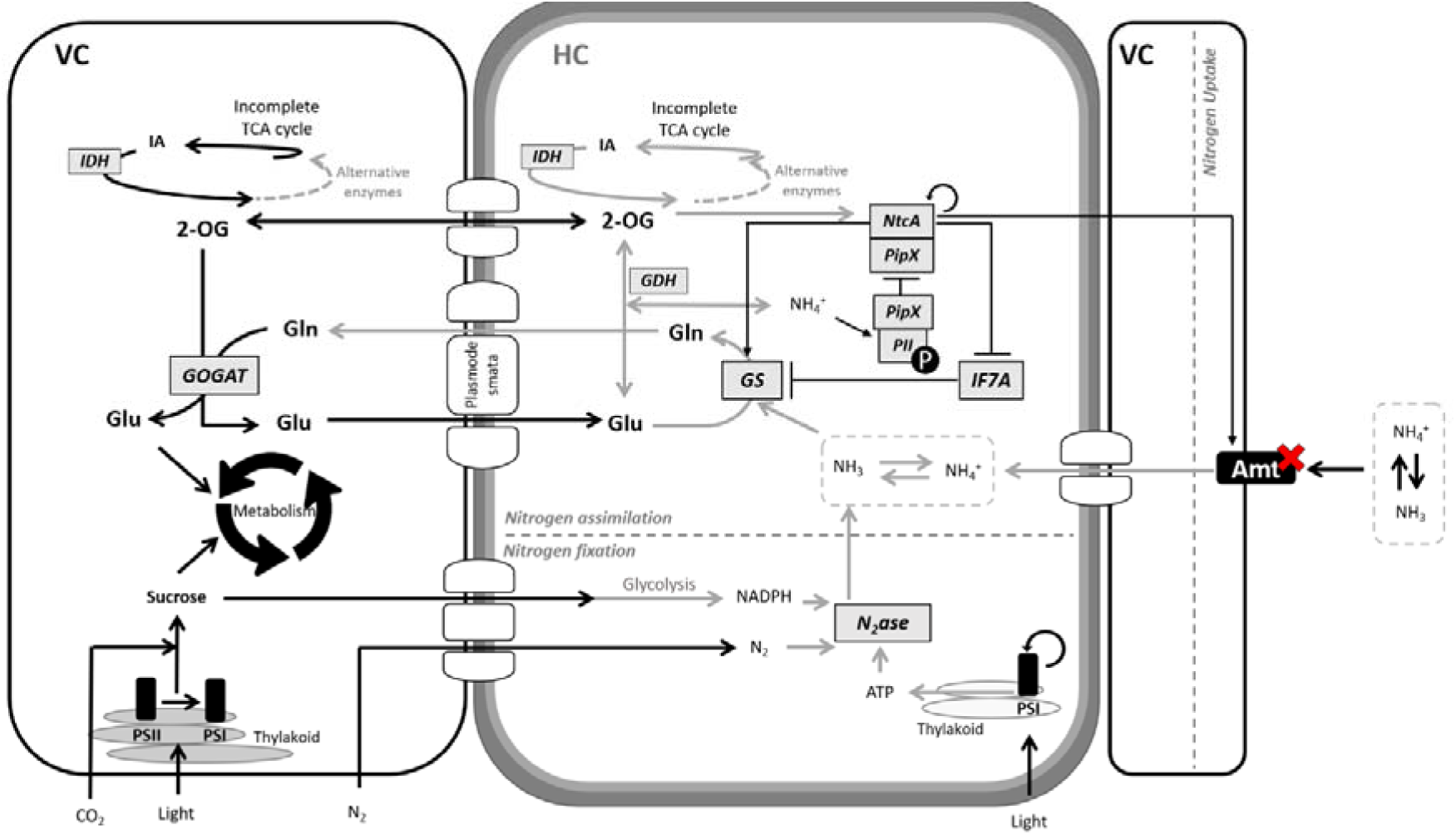
Schematic overview of major molecular players regulating N metabolism and the metabolic interaction between heterocysts (HC) and vegetative (VC) cells in Anabaena sp. PCC 7120, in diazotrophic conditions. Nitrogenase (N_2_ase) fixes one molecule of atmospheric N_2_ into two molecules of NH_3_ in heterocysts (Inomura et al, 2017), using reducing power (NADPH) from the catabolism of carbon -compounds (sucrose) photosynthesised in vegetative cells (Cumino et al, 2007; Nürnberg et al, 2015) and energy (ATP) from the residual photosynthetic activity in heterocysts [i.e. cyclic electron flow around photosystem I (PSI), (Cardona & Magnuson, 2010)]. NH_3_ is then assimilated through glutamine synthetase (GS) via the amidation of glutamate (Glu) to glutamine (Gln) (Forchhammer & Selim, 2019). GS activity is controlled through posttranslational inactivation (Bolay et al, 2018) by IF7A (Galmozzi et al, 2010). Glutamate dehydrogenase (GDH) marginally contributes to the assimilation flux of fixed N, catalysing the reversible conversion of 2-oxoglutarate (2-OG) to Glu. Subsequently, in vegetative cells (Martín-Figueroa et al, 2000), glutamine oxoglutarate aminotransferase (GOGAT) catalyses the transfer of the amine group from Gln to 2-OG, generating two molecules of Glu. As N metabolism spans different cell types, a coordinated exchange of metabolites (i.e. sucrose, Gln, Glu and 2-OG) between vegetative cells and heterocysts via septal junctions [plasmodesmata (Mullineaux et al, 2008)] is required to maintain metabolic homeostasis. 2-OG is also a metabolic intermediate of the tricarboxylic acid (TCA) cycle [synthesised from isocitric acid (IA) by isocitrate dehydrogenase (IDH)], thus connecting N and C metabolism at a central point. N metabolism homeostasis is controlled by a molecular network, including the proteins NtcA, PipX and PII. When external N is available, PII is not phosphorylated and it sequesters PipX, preventing its biding to NtcA and consequently its activation. When N is limiting, PII is phosphorylated, freeing PipX, which ultimately binds and activates NtcA (Flores & Herrero, 2005). The red cross indicates the knock-out (KO) mutant Δamt used in this work (Paz-Yepes et al, 2008). Major molecular players targeted in this work are highlighted by a grey square or in bold, respectively for proteins and metabolites.

Heterocysts and vegetative cells have complementary metabolism, with the former providing fixed nitrogen and the latter returning reduced carbon needed to sustain BNF (Malatinszky *et al*, 2017). This metabolic exchange and associated networks (summarised in Fig. 1) are most likely carefully coordinated in order to ensure organism-level homeostasis (Mullineaux *et al*, 2008). The question is, how does this molecular coordination take place?

In many organisms, including 7120, N metabolism is orchestrated by a complex signalling network with the likely aim to balance the cellular C/N ratio (Forchhammer & Selim, 2019). N and C metabolism are in fact tightly coupled as (1) the two elements are the most abundant in living organisms, calling for coordination to avoid metabolic inefficiencies, and (2) N assimilation depends on the availability of C skeleton, with shortage or oversupply strongly affecting the metabolism of N (Zhang *et al*, 2018). Therefore, a properly balanced N and C metabolism is necessary for optimal growth and different levels of regulation exist to control uptake and assimilation efficiencies of both chemical species. When the C source (i.e. CO_2_ in case of phototrophic metabolism) is not limiting, the regulatory mechanisms controlling C/N balance depend on both the abundance and the nature of the N sources available to the cell. Although cyanobacteria can use multiple N sources, including NH_4_^+^, intracellularly they are all converted to NH_4_^+^, the most reduced and energetically favourable N source (Robinson, 2017). Ammonia translocation across biological membranes is actively driven by AMT transporters that belong to a family of permeases widely distributed in living organisms (Javelle *et al*, 2007), or through passive diffusion if the external pH pushes the equilibrium towards the uncharged form (NH_3_, ammonia). 7120 bears a gene cluster including three *amt* genes, namely *amt4, amt1* and *amtB* (Paz-Yepes *et al*, 2008). In this work, we used a knock-out (KO) mutant of the whole gene cluster in 7120 [henceforth Δ*amt*, (Paz-Yepes *et al*, 2008)] to perturb both the sensing of external N and low-affinity uptake of NH4^+^, with the aim to investigate how a N_2_-fixing cyanobacterium responds to perturbation of N-metabolism at a whole-system level. Although several genes and proteins in 7120 have been individually studied previously (Flores & Herrero, 2005; Forchhammer & Selim, 2019), it is difficult to make over-arching conclusions on the regulatory system, also as cyanobacteria differ substantially relative to heterotrophic bacteria (Reitzer, 2003; Bolay *et al*, 2018). The aim of this work is also to enhance our understanding that contributes towards the practical goal of eventually rerouting N -metabolism for biotechnological purposes (Perin *et al*, 2019).

## Results

### Amt transporters are not required to support growth of 7120 under constant laboratory conditions but play a role in N metabolism

Phototrophic growth of 7120 WT and Δ*amt* strains was carried out in media with different N sources (NO_3_^-^, N_2_ and NH_4_^+^) at saturating CO_2_ (i.e. 1%), in order to avoid C limitation. Such conditions are expected to vary the internal C/N balance by modifying the abundance and nature of the N source. The Δ*amt* mutant did not display any growth phenotype with respect to the parental strain, regardless of the N source (Fig. 2A), confirming that the whole *amt* cluster is not necessary to support growth of 7120 under the tested laboratory conditions (Paz-Yepes *et al*, 2008).

**Figure 2.**
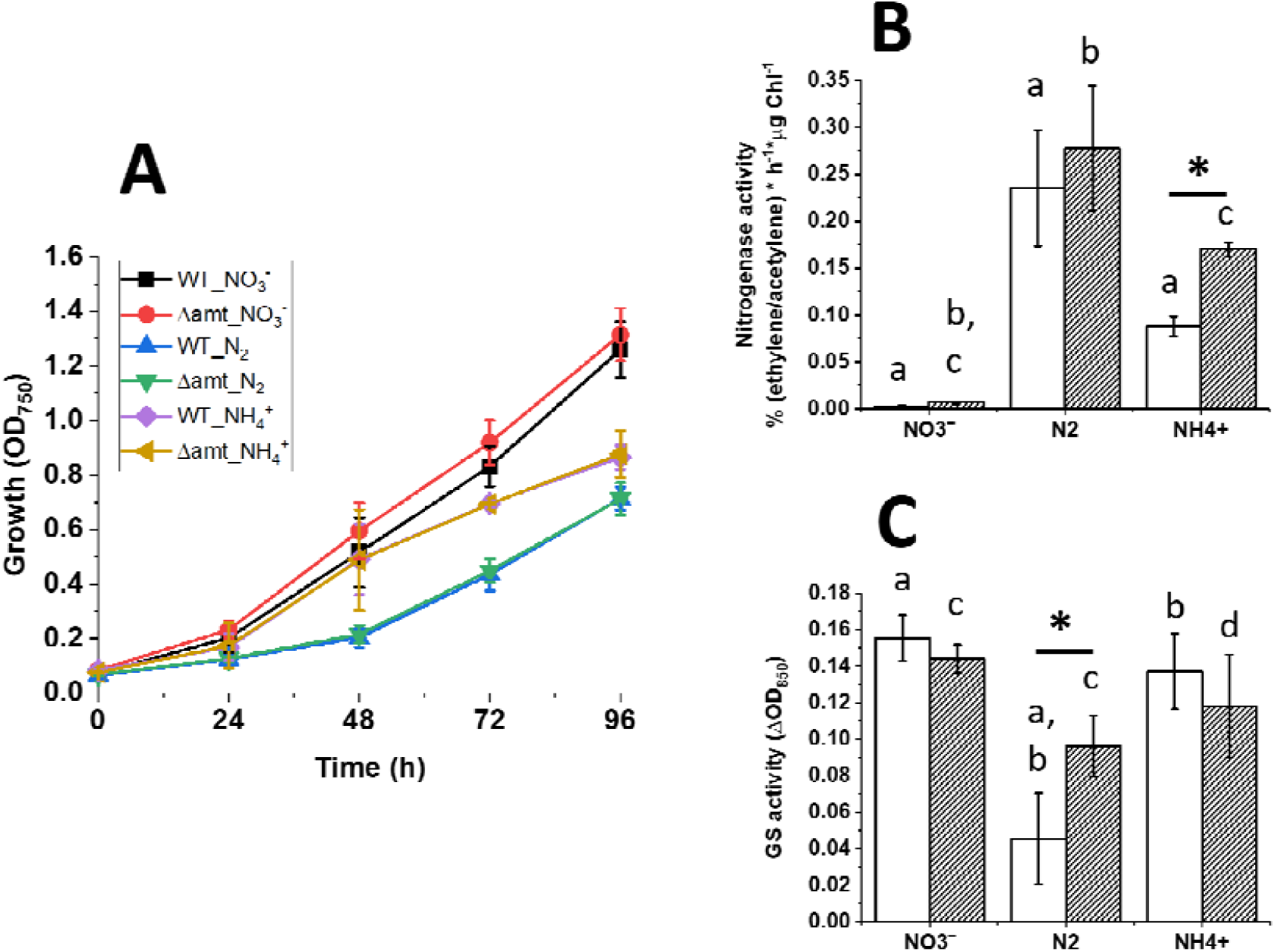
Growth of 7120 WT and Δamt strains with different N sources. The two strains were cultivated in different N sources for 96 h. Growth (A) was monitored over the course of the whole experiment, whilst Nitrogenase activity (B) and GS activity (C) were measured after 96 h in such cultivation conditions. Data are indicated as average ± SD of 6 biological replicates. Statistically significant differences between WT (white bars) and Δamt (striped bars) are indicated with an asterisk, whilst the same alphabet letter indicates statistically significant differences for the same strain in different growth conditions (one-way ANOVA, p-value < 0.05).

Growth in atmospheric N_2_ lags behind both NO_3_ ^-^ and NH_4_ ^+^, with cells taking ∼48 hours to fully switch to atmospheric N_2_ fixation. Growth in NH_4_^+^ shows two distinct modes, as indicated by the specific growth rate (Supplementary Fig. S2), suggesting that 5 mM NH_4_^+^ is not enough to support maximal growth (i.e. growth in NO_3_^-^ in this experiment) over the whole experimental time frame, and that it likely runs out after ∼48 hours (Fig. 2A). The nitrogenase activity, measured after 96 hours, confirms that 5 mM NH_4_ ^+^ runs out over the course of the experiment, triggering diazotrophic growth (Fig. 2B). Moreover, the lower nitrogenase activity with respect to N_2_ conditions highlights that cells in NH4 ^+^ media, at the time of sampling, are in a transitory phase before reaching the maximal N fixation potential. Interestingly, in NH4^+^ media, the Δ*amt* strain shows a higher nitrogenase activity per unit of chlorophyll (Chl) than the parental strain, although that does not result in an improvement in growth, suggesting possible compensatory modifications in the following metabolic steps (e.g. N assimilation). After 96 h, GS activity, a (supposed) central player in N metabolism in this organism (Bolay *et al*, 2018), is also affected by the mutation (as in the case of fully diazotrophic conditions (N_2_) in which Δ*amt* strain shows a greater N assimilation activity than the WT, see Fig. 2C). Moreover, when both strains are grown in N_2_, GS activity is overall lower than in the other two growth conditions. The collective data indicated that the loss of AMT resulted in no phenotypic change, but that N-metabolism had adjusted, presumably to maintain homeostasis, raising the following question: how extensive was this adaption and what molecular players were involved?

In 7120, GS is regulated both transcriptionally and post-translationally, according to the C/N status of the cell (Bolay *et al*, 2018). The abundance of the protein is transcriptionally controlled and changes according to the N source(s) as observed in the WT strain, but not in Δ*amt* (Fig. 3A).

**Figure 3.**
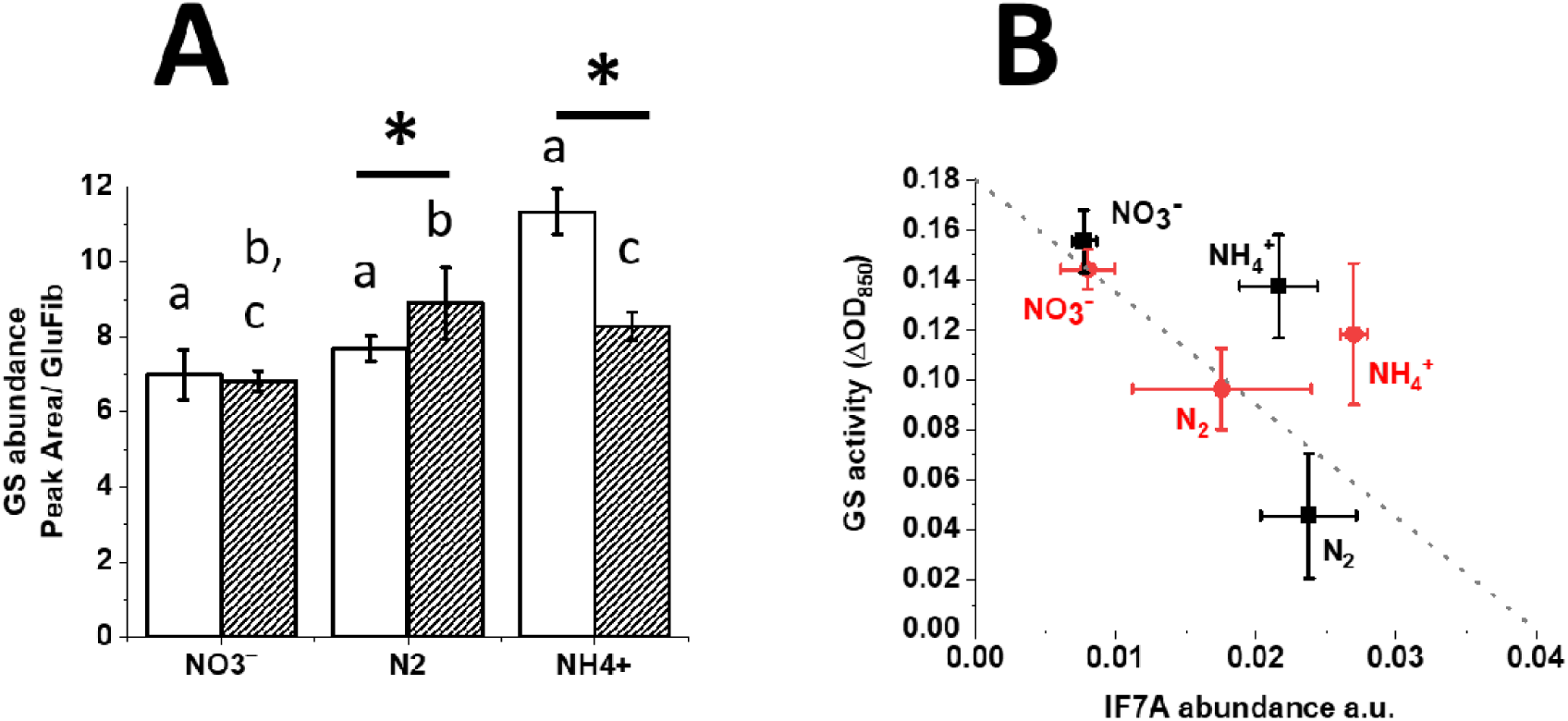
GS abundance and correlation between GS activity and IF7A amount for 7120 WT and Δ*amt* strains, after 96 h in the growth conditions of figure 2A. IF7A quantification data are reported in supplementary Fig. S3. Data are indicated as average ± SD of 6 biological replicates. *Statistically significant differences between WT (white bars and black squares) and* Δ*amt (striped bars and red circles) are indicated with an asterisk, whilst the same alphabet letter indicates statistically significant differences for the same strain in different growth conditions (one-way ANOVA, p-value < 0.05)*.

The abundance of GS (Fig. 3A) does not reflect its measured activity (Fig. 2C). GS activity is known to be controlled by covalent binding of the inactivation factor IF7A, in response to the C/N balance of the cell (Galmozzi *et al*, 2010). As shown in Fig. 3B, in the WT strain, GS activity broadly displays a negative correlation with the amount of the inactivation factor IF7A, as expected. There is no linear correlation for either of the strains across different N sources, however, suggesting other molecular players also contribute to the regulation of GS activity in this organism. The absence of AMT transporters has an effect on the IF7A/GS activity relationship (Fig. 3B). In N replete conditions (NO_3_ ^-^), there is no difference between the two strains, whilst under the two other N deplete conditions, the deletion of *amt* results in a divergence between the two strains (Fig. 3B). The genetic and environmental treatments affect GS activity through a combined variation in both GS and IF7A abundance (Fig. 3A and supplementary Fig. S3). In particular, it is worth noting that Δ*amt* retains GS activity even if the amount of IF7A changes, supporting the idea that other regulatory players also may be involved. Based on the available data, we hypothesised that AMT transporters are directly or indirectly involved in the regulation of GS activity during the switch towards diazotrophic conditions in 7120, as already observed in the purple bacterium *Rhodobacter capsulatus* (Yakunin & Hallenbeck, 2002).

### Amt transporters regulate N metabolism homeostasis in 7120

In order to test this hypothesis, the internal metabolic changes taking place in 7120 WT and Δ*amt* during the transition from N replete to deplete conditions were followed by combining physiological data with targeted proteomic and metabolomic quantification of molecular players known to be involved in the regulation of N metabolism in 7120 (Fig. 1). Both WT and Δ*amt* strains were cultivated in BG11_0_ supplemented with 5 mM NH_4_^+^. The concentration of ammonium/ammonia in the media and cell growth was monitored over time (Fig. 4A). Four different time points were chosen to investigate the physiological and metabolic status of the cells [i.e. NR (N replete conditions), ND1, ND2 and ND4 (1, 2 and 4 days, respectively, after N depletion), corresponding to 24, 72, 96 and 144 h from the start of the experiment]. Former studies in 7120 mainly focused on the first 24 h after N deprivation (Galmozzi *et al*, 2010; Valladares *et al*, 2011), as heterocyst differentiation is expected to occur within such time frame (Valladares *et al*, 2011). Here, instead, we opted for an extended sampling protocol in order to complement the information already available in literature with the knowledge of the metabolic/proteomic adjustments happening over a longer time-scale.

**Figure 4.**
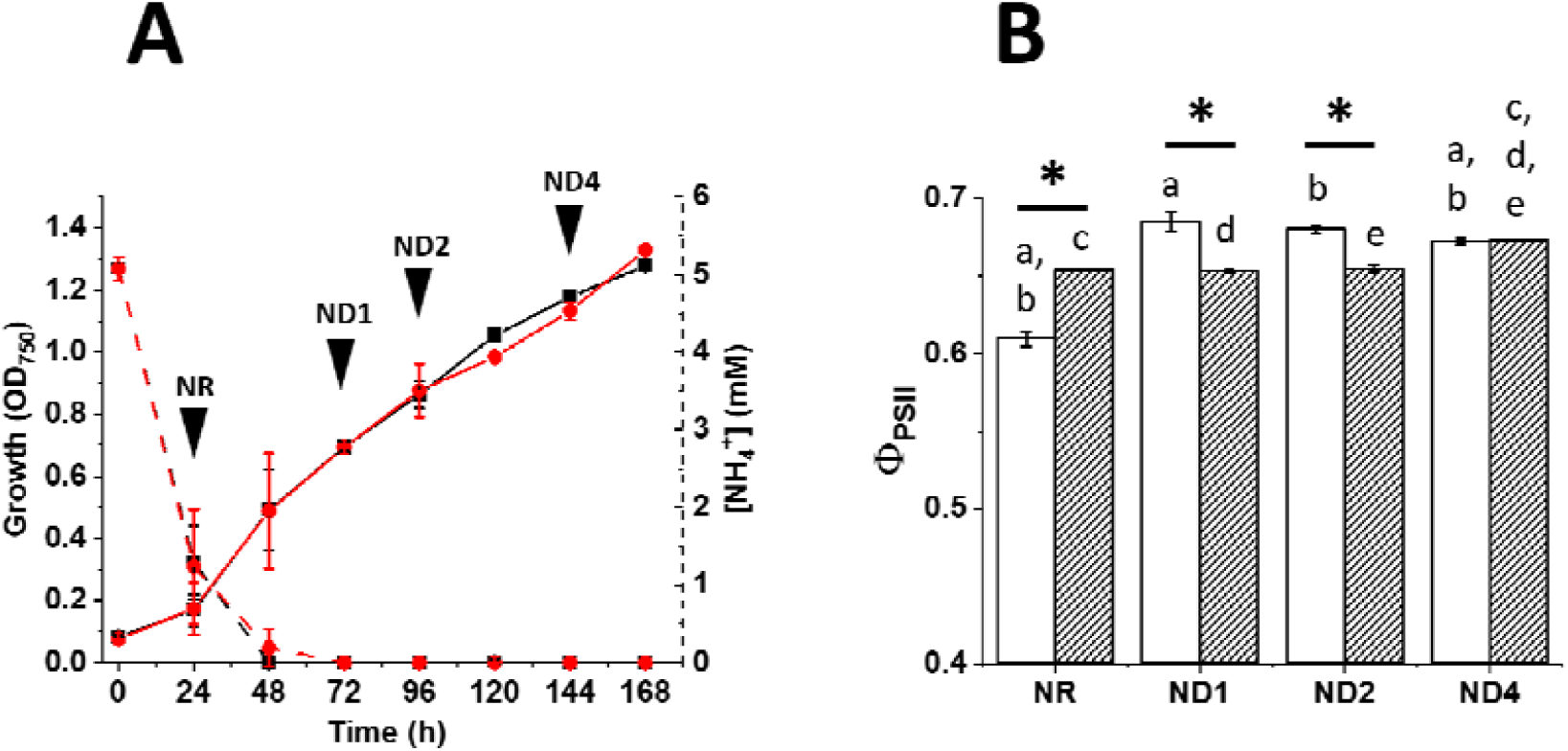
Growth, ammonium/ammonia consumption (A) and maximal photosynthetic efficiency [(Φ_PSII_), (B)] monitoring for 7120 WT and Δ*amt* strains in BG11_0_ + 5 mM NH_4_^+^. In A., black and red dashed lines indicate ammonium/ammonia concentration over time for 7120 WT and Δ*amt* strains, respectively. Cultures were sampled at four time points over the course of the experiment [NR (Nitrogen Replete), ND1, ND2 and ND4, respectively 1, 2 and 4 days after Nitrogen Deprivation]. Data are indicated as average ± SD of 6 biological replicates. Statistically significant differences between WT (black squares and white bars) and Δ*amt* (red circles and striped bars) are indicated with an asterisk, whilst the same alphabet letter indicates statistically significant differences for the same strain in different growth conditions (one-way ANOVA, p-value < 0.05).

The absence of the whole *amt* cluster does not affect the ammonium/ammonia consumption rate, indicating the diffusion of ammonia is enough to sustain growth in 7120, under the tested experimental conditions (Fig. 4A). Moreover, ammonium/ammonia in the medium is fully depleted after 48 h (Fig. 4A), confirming 5 mM NH_4_^+^ is not enough to support maximal growth in 7120 over a 96 h-long experiment (Fig. 2A). Over the course of the experiment, both strains mostly showed a stable pigment content, suggesting the switch towards diazotrophic conditions does not unbalance the overall N status of the cell (Table 1).

**Table 1.** Pigment content of WT and Δ*amt* strains during the switch from N replete to deplete conditions. Cultures were sampled at four time points over the course of the experiment [NR (Nitrogen Replete), ND1, ND2 and ND4, respectively 1, 2 and 4 days after Nitrogen Deprivation], according to Fig. 4A. Data are indicated as average ± SD of 6 biological replicates. Statistically significant differences between WT and Δ*amt* are indicated with an asterisk, whilst the same alphabet letter indicates statistically significant differences for the same strain in different growth conditions (one-way ANOVA, p-value < 0.05).

| Chl content<br>($\mu\text{g}/\text{OD}_{750}$) | WT | Δamt |
| --- | --- | --- |
| NR | 13.36 ± 0.49<sup>a</sup> | 14.14 ± 0.55<sup>d</sup> |
| ND1 | 13.41 ± 0.18<sup>b</sup> | 13.38 ± 0.41<sup>e</sup> |
| ND2 | 12.37 ± 0.37<sup>*,a,b,c</sup> | 14.26 ± 0.27<sup>*,e</sup> |
| ND4 | 14.49 ± 0.72<sup>*,c</sup> | 16.13 ± 0.41<sup>*,d,e</sup> |
| Car content<br>($\mu\text{g}/\text{OD}_{750}$) | | |
| NR | 5.78 ± 0.2<sup>*,a</sup> | 4.73 ± 0.22<sup>*,c</sup> |
| ND1 | 4.84 ± 0.05<sup>a,b</sup> | 4.7 ± 0.13<sup>d</sup> |
| ND2 | 4.49 ± 0.12<sup>a,b</sup> | 4.76 ± 0.11<sup>e</sup> |
| ND4 | 5.54 ± 0.29<sup>b</sup> | 5.86 ± 0.18<sup>c,d,e</sup> |

| Chl/Car |  |  |
| --- | --- | --- |
| NR | 2.31 ± 0.005<sup>*,a</sup> | 2.99 ± 0.02<sup>*,c</sup> |
| ND1 | 2.77 ± 0.01<sup>*,a</sup> | 2.85 ± 0.01<sup>*,c,d</sup> |
| ND2 | 2.75 ± 0.01<sup>*,b</sup> | 2.99 ± 0.01<sup>*,d</sup> |
| ND4 | 2.61 ± 0.009<sup>*,a,b</sup> | 2.75 ± 0.01<sup>*,c,d</sup> |

However, the Δ*amt* strain shows a greater Chl/Car ratio, mainly achieved through the accumulation of a higher Chl content than the parental strain (Table 1). This effect on the pigment composition has also a consequence on the photosynthetic performances. The photosynthetic activity in the parental strain changes over the course of the experiment, whilst in Δ*amt* it is more stable (Fig. 4B). Moreover, the mutant shows a higher photosynthetic efficiency than WT in N replete conditions, while the difference reverses both after 24 and 48 h under N deprivation conditions and ultimately disappears after 96 h (Fig. 4B). Given that the Δ*amt* mutation does not trigger any growth phenotype, whilst there are substantial changes to both photosynthesis and N metabolism, we hypothesised that the deletion of the whole *amt* cluster is in fact triggering a whole cell metabolic response in order to maintain homeostasis.

In order to validate the two unanswered hypotheses, we investigated the N metabolism of 7120 more closely, targeting the same proteins and enzymatic reactions as above, but this time measured in the same time intervals as indicated in Fig. 4A. Both WT and Δ*amt* strains activate N fixation (Fig. 5A) as a consequence of ammonium/ammonia deprivation (Fig. 4A). The Δ*amt* strain shows a higher nitrogenase activity than the parental strain in both ND1 and ND2, indicating a faster response to N deprivation than the parental strain (Fig. 5A). Abundance of NifK and NifD, encoding for α and β subunits of nitrogenase, is higher in the mutant strain (Fig. 5B and 5C, respectively), suggesting increased N fixation activity depends at least in part on a greater accumulation of the protein complex, given also the number of heterocysts over the course of the experiment is not affected by the mutation (Supplementary Fig. S4). The increased N fixation activity does not translate to a greater growth rate in the mutant, however, suggesting possible compensatory modifications in downstream steps of N metabolism.

**Figure 5.**
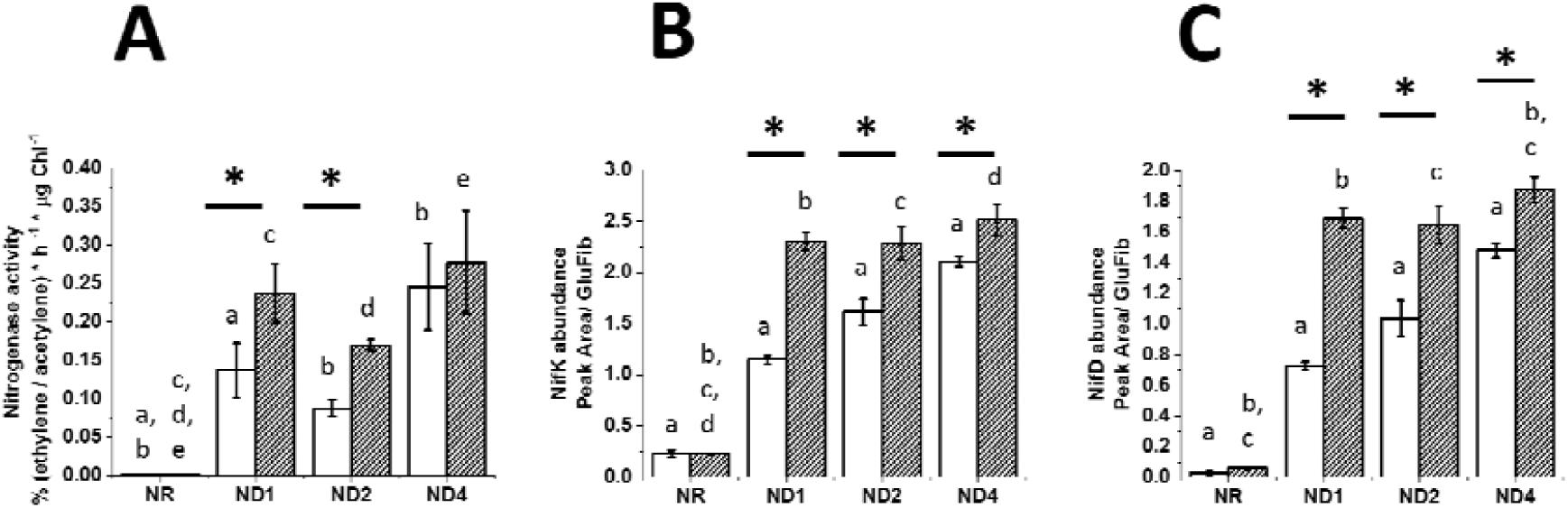
Nitrogenase activity (A), NifK (B) and NifD (C) abundance in both WT and Δamt strains, following N deprivation. Time points are those indicated in Fig. 4A and correspond to NR (Nitrogen Replete), ND1, ND2 and ND4, respectively 1, 2 and 4 days after Nitrogen Deprivation. Data are indicated as average ± SD of 6 biological replicates. Statistically significant differences between WT (white bars) and Δamt (striped bars) are indicated with an asterisk, whilst the same alphabet letter indicates statistically significant differences for the same strain in different growth conditions (one-way ANOVA, p-value < 0.05).

Once atmospheric N is fixed into ammonium/ammonia, the latter is incorporated into amino acid metabolism. GS activity was strongly regulated in both strains also in this experiment, as observed before (Fig. 2B). In N replete conditions, Δ*amt* showed greater N assimilation activity than the parental strain (i.e. NR in Fig. 6A), likely to be the cause of the increased influx of N in the central metabolism, as indicated by the increased pigment content and photosynthetic activity observed in NR conditions (Table 1 and Fig. 4B). Consequently, GS activity was influenced by the change in N metabolism, especially in the earlier samples. GS activity was at first reduced and then strongly increased (i.e. ND1 and ND2 in fig. 6A), before stabilising again to the same rate observed under N replete conditions (i.e. ND4 in fig. 6A). The trend in GS activity appeared to depend on the abundance of both GS and IF7A (Fig. 6B and 6C, respectively), which accumulate differentially in the two strains. The deletion of the whole *amt* cluster in fact strongly affects the abundance of both proteins, which impacts the overall regulation of GS activity, possibly calling for other compensatory mechanisms in this genetic background, as often GS activity is retained even though differences in the amount of IF7A are present (Fig. 6D). These results strengthen the hypothesis that AMT transporters play a direct or indirect IF7A-independent role on the GS activity in 7120.

**Figure 6.**
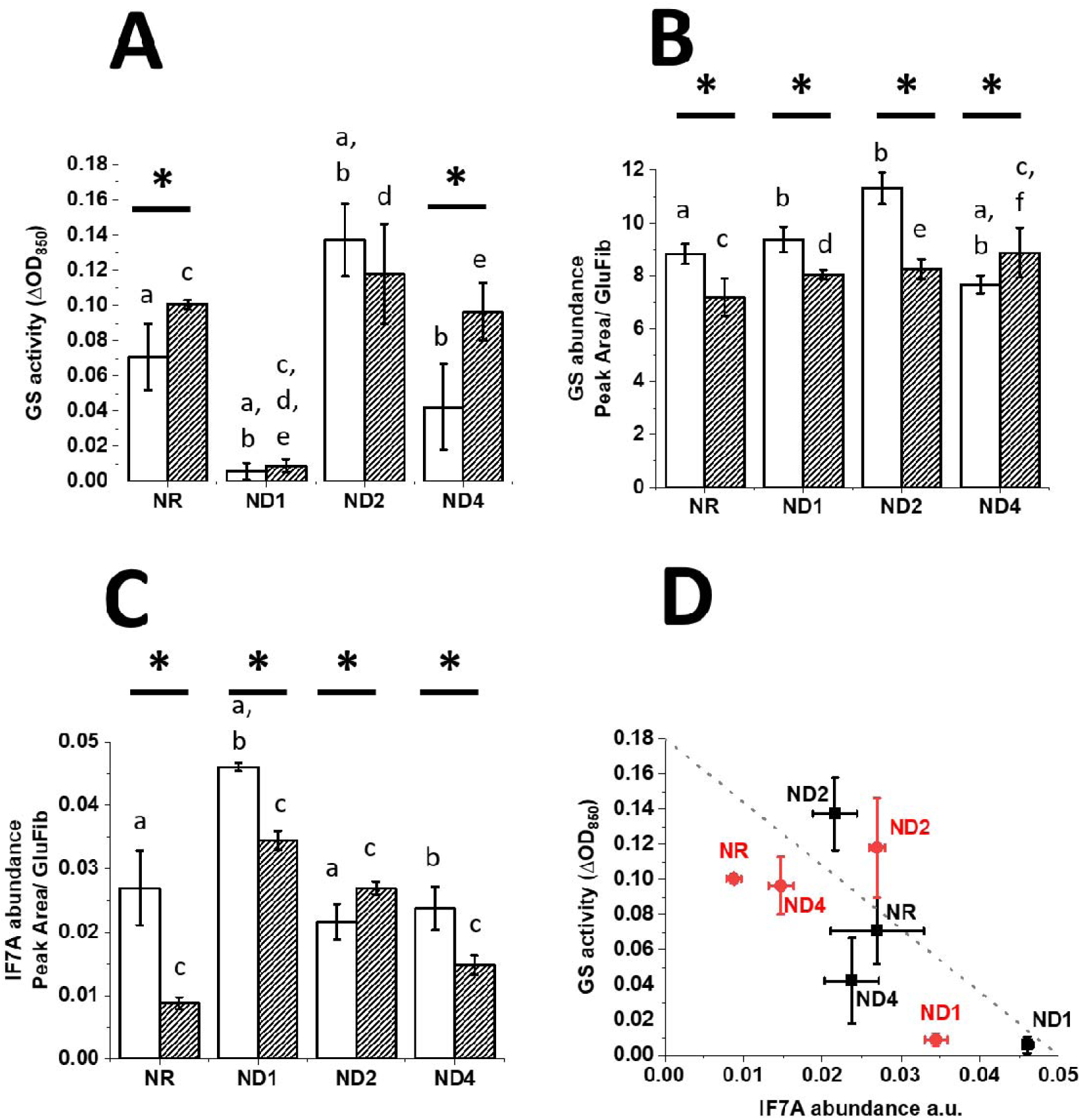
Regulation of N assimilation during the switch towards N deprivation in both 7120 WT and Δ*amt* strains. A. GS activity; B. GS abundance; C. IF7A abundance; D. Correlation between GS activity and IF7A abundance. Time points are those indicated in Fig. 4A and correspond to NR (Nitrogen Replete), ND1, ND2 and ND4, respectively 1, 2 and 4 days after Nitrogen Deprivation. Data are indicated as average ± SD of 6 biological replicates. Statistically significant differences between WT (white bars and black squares) and Δ*amt* (striped bars and red circles) are indicated with an asterisk, whilst the same alphabet letter indicates statistically significant differences for the same strain in different growth conditions (one-way ANOVA, p-value < 0.05).

GS is the major entry point of fixed N in the central metabolism of 7120 (Bolay *et al*, 2018). Nevertheless, other enzymes control the availability of GS substrates, thus indirectly contributing to the regulation of N assimilation. These include GOGAT (responsible for the regeneration of Glu in the GS-GOGAT cycle), IDH (responsible for the synthesis of 2-OG, a substrate of GOGAT) and GDH (involved in the reversible conversion between Glu and 2-OG) (Martín-Figueroa *et al*, 2000; Bolay *et al*, 2018; Forchhammer & Selim, 2019). Under the tested experimental conditions, the deletion of the whole *amt* cluster had major consequences also on the abundance of such enzymes (Supplementary fig. S5). GOGAT accumulation followed the same trend in both strains over the course of the experiment (i.e. strong downregulation as a consequence of N deprivation, supplementary fig. S5A), while IDH and GDH displayed different trends in the two genetic backgrounds (Supplementary fig. S5B and S5C, respectively). These observations support the notion that the absence of AMT transporters triggers both direct and indirect effects on N metabolism in 7120.

### The absence of AMT transporters affects the master regulatory network of N metabolism

Given the impact of Δ*amt* on both GS and nitrogenase, the question is how widespread the adjustments had rippled further into the cellular system? In order to address this question, we investigated the N metabolism more deeply, with an expanded number of protein quantification targets. The C/N balance of the cell is in fact also known to regulate the interaction between NctA, PipX and PII in 7120 (Forchhammer & Selim, 2019), which are expected to be the major molecular players controlling the metabolic remodelling in response to both the nature and availability of N source in the external environment, in 7120 (Fig. 1). The transcription factor NtcA, active only once bound to PipX, regulates the abundance of nitrogenase, GS and IF7A (Flores & Herrero, 2005). As Δ*amt* both responds to N deprivation more quickly than the parental strain, by activating faster N fixation (Fig. 5), and also shows major alterations in the regulation of N assimilation (greater GS activity), we wondered whether the mutation might have an effect on such master molecular regulators, which control both enzymatic steps in this species (i.e. NtcA, PII and PipX, Fig. 1) and we thus quantified their abundance (Fig. 7).

**Figure 7.**
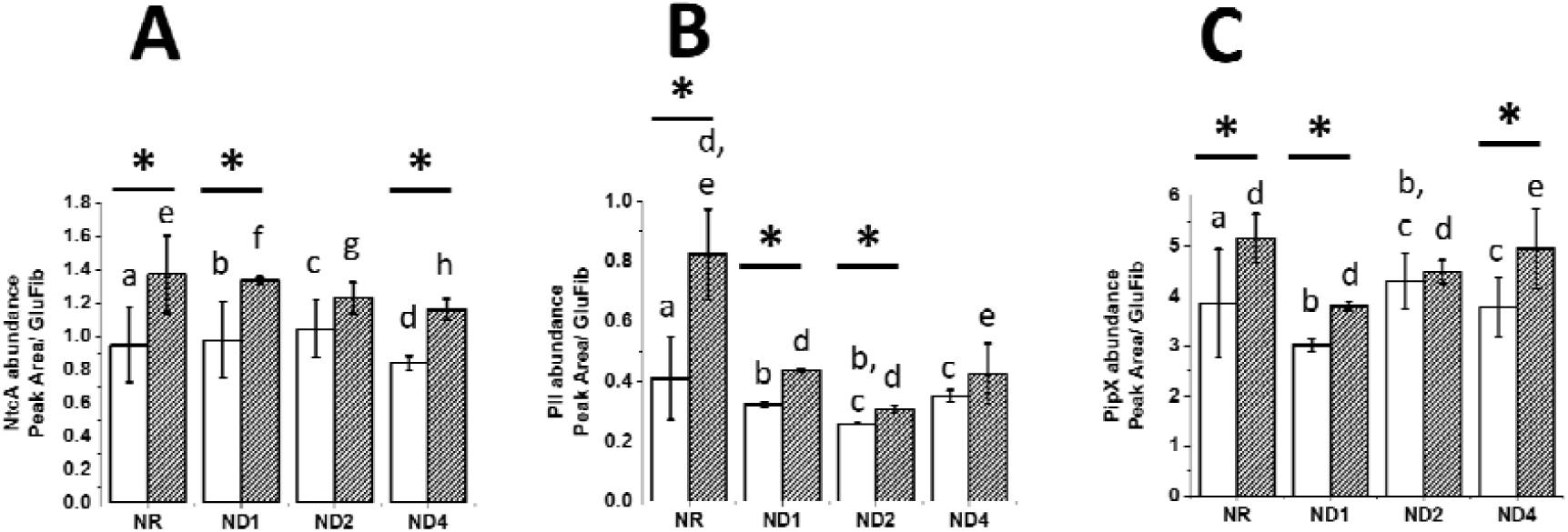
Abundance of the three major molecular players regulating N metabolism in 7120. A. NtcA; B. PII; C. PipX. Time points are those indicated in Fig. 4A and correspond to NR (Nitrogen Replete), ND1, ND2 and ND4, respectively 1, 2 and 4 days after Nitrogen Deprivation. Data are indicated as average ± SD of 6 biological replicates. Statistically significant differences between WT (white bars) and Δ*amt* (striped bars) are indicated with an asterisk, whilst the same alphabet letter indicates statistically significant differences for the same strain in different growth conditions (one-way ANOVA, p-value < 0.05).

Overall, Δ*amt* accumulated more of the three proteins over the course of the experiment relative to the parental strain, suggesting the absence of AMT transporters has an extensive impact on the cellular system, involving the master regulatory network of N metabolism. This might explain the more rapid activation of N -fixation in response to N deprivation and also the effect on N assimilation, discussed above (Fig. 5 and Fig. 6). Moreover, while the abundance of the three proteins does not vary much over the course of the experiment in WT, PII does respond to N deprivation and is more abundant under NR conditions in Δ*amt* (Fig. 7B). This suggests PII might be degraded after induction of N deprivation, possibly as a consequence of a greater phosphorylation rate (see Fig. 1 for the molecular mechanisms controlling PII/PipX interaction). These results strengthen the notion that AMT transporters play a central role in the regulation of N metabolism in 7120.

### Metabolic remodelling as a consequence of amt deletion

Under diazotrophic conditions, coordinated metabolic interaction between vegetative cells and heterocysts is seminal for optimal growth. The absence of AMT transporters triggered an extensive remodelling at the protein level in 7120, spanning both cell types, given some of the proteins investigated in this work are known to be exclusively expressed in one of the two cell types [e.g. GOGAT in vegetative cells, (Martín-Figueroa *et al*, 2000)]. We therefore wondered whether the same also happens at the metabolite pool level. Cyanobacteria evolved the ability to store assimilated N in the form of cyanophycin granule polypeptide (CPG), possibly acting as a buffer to naturally varying N -fixation due to fluctuations in N supply and day/night cycles (Watzer & Forchhammer, 2018). In heterocystous filamentous cyanobacteria, CPG accumulates at the contact sites between heterocysts and adjacent vegetative cells and is expected to regulate the transfer of fixed N from the former to the latter (Burnat *et al*, 2014), thus influencing metabolite exchange between the two cell types (see supplementary Fig. S6A for a schematic overview of CPG metabolism in 7120). In order to investigate whether the metabolites exchange between the two cell types was also affected by the mutation, we studied potential alterations in the CPG metabolism (Fig. 8).

**Figure 8.**
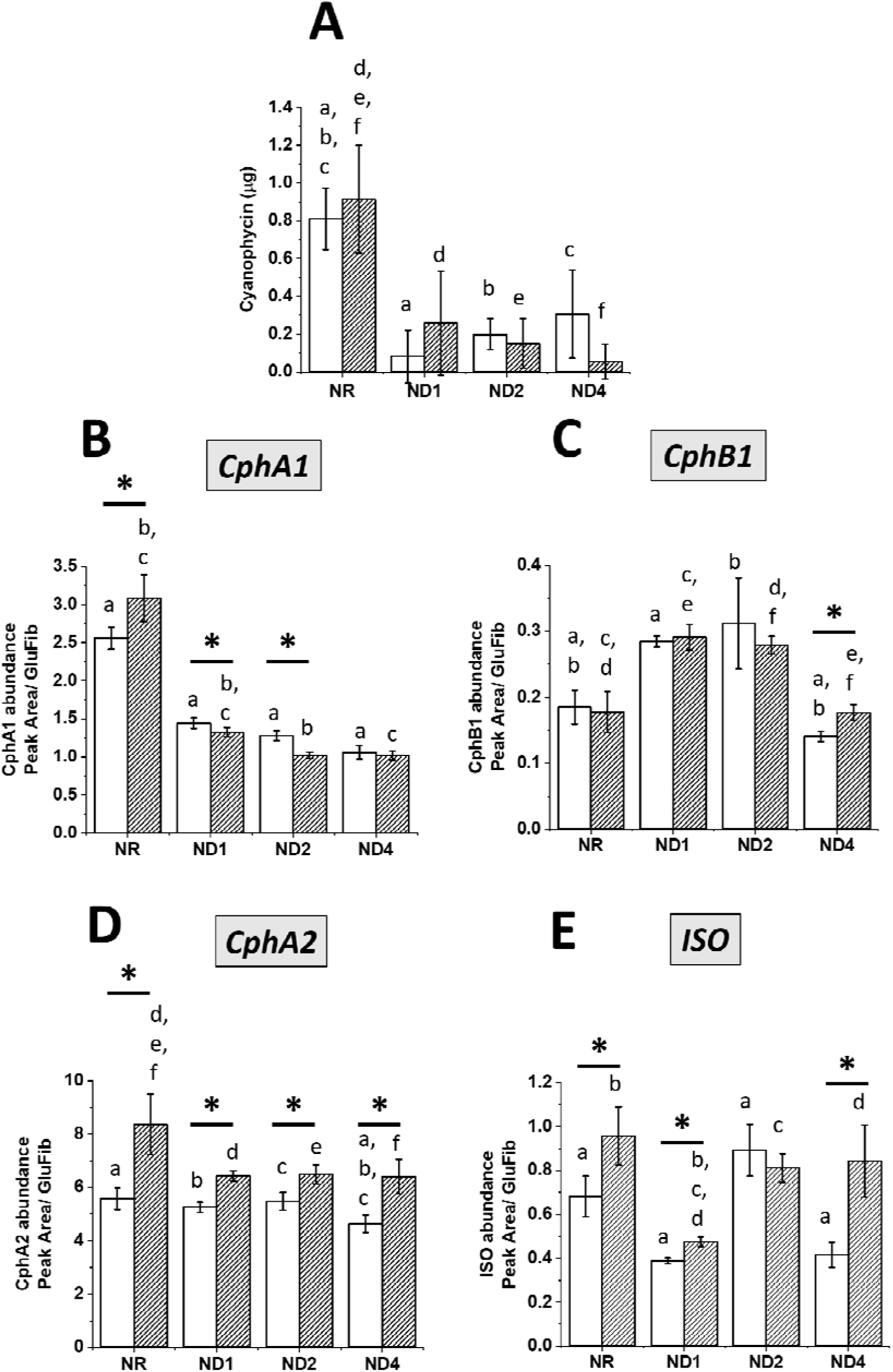
Cyanophycin (CPG) content and abundance of the four major enzymes regulating its metabolism in both WT and Δamt strain. A. Cyanophycin content in both WT and Δamt strains in the four time points chosen in this experiment, according to Fig. 4A. The same amount of biomass was processed in all conditions for both strains (see materials and methods). B, C, D and E. Abundance of the four major proteins [i.e. Cyanophycin synthetase (CphA1), Cynaophycinase (CphB1), Cyanophycin synthetase 2 (CphA2) and Isoaspartyl dipeptidase (ISO), respectively for B, C, D and E] regulating cyanophycin metabolism in 7120, in the four time points chosen in this experiment (Fig. 4A). Data are indicated as average ± SD of 6 biological replicates. Statistically significant differences between WT (white bars) and Δamt (striped bars) are indicated with an asterisk, whilst the same alphabet letter indicates statistically significant differences for the same strain in different growth conditions (one-way ANOVA, p-value < 0.05). See supplementary fig. S6 for an overview of CPG metabolism.

Under the tested experimental conditions, both strains accumulated CPG in NR conditions as expected (Forchhammer & Watzer, 2016), suggesting the deletion of AMT transporters did not perturb N storage to the point of affecting CPG accumulation. In fact, both strains show the same CPG content in NR (Fig. 8A), indicating the pool of available CPG to support the metabolic needs of the cell is not affected by the mutation at this time point. Nevertheless, the abundance of three out of four major enzymes controlling CPG metabolism (Supplementary Fig. S6) is significantly different between the two strains, with CphA1, CphA2 and ISO showing increased accumulation in the mutant with respect to the parental strain in NR (Fig. 8B, D and E). We therefore hypothesised that the overall CGP metabolic flux is accelerated in the mutant under NR conditions, possibly enabling a faster response to environmental changes. When cells experience N deprivation, the cyanophycin content decreases, presumably as it is rapidly used as N source (Forchhammer & Watzer, 2016) (i.e. after one day of N deprivation CPG is fully consumed, Fig. 8A). Nevertheless, whilst CPG starts building up again in the parental strain after two days of N deprivation, a constant consumption trend is observed over the course of the experiment in the mutant (Fig. 8A). This suggests that N fixation in the WT exceeds metabolic needs and a fraction of the assimilated N is thus stored as CPG, while in the mutant this trend is disrupted. Out of the four major enzymatic steps controlling CPG metabolism in 7120, cyanophycin synthetase (CphA1), the major enzyme controlling CPG biosynthesis (Forchhammer & Watzer, 2016), is most affected by N deprivation (Fig. 8 B, C, D and E), with the mutant showing a faster reduction in its abundance than the parental strain (Fig. 8B). It is also worth noting that cyanophycin synthetase 2 (CphA2), a truncated version of CphA1 catalysing the direct recycling of the β-aspartyl-arginine dipeptide into CPG (Supplementary Fig. S6, (Forchhammer & Watzer, 2016)], is more abundant in the mutant strain in all time points (Fig. 8D), strengthening the notion that CPG metabolism is accelerated in Δ*amt*.

CPG accumulation in N fixing cyanobacteria is mediated by PII which in turn regulates N-acetyl-N-glutamate kinase (NAGK) activity (Forchhammer & Selim, 2019). NAGK catalyses the conversion of N-acetyl-L-glutamate to N-acetyl-L-glutamyl-phosphate, which is further converted to ornithine, from where Arg, the end -product of the pathway, is derived. Therefore, CPG biosynthesis directly follows the concentration of free Arg in the cell, as a consequence of feedback inhibition of NAGK (Watzer *et al*, 2015). In our experimental conditions, the free Arg concentration indeed strongly decreased upon N deprivation in both strains (Fig. 9A and B), confirming the strong reduction in CPG content upon N deprivation, observed before (Fig. 8A).

**Figure 9.**
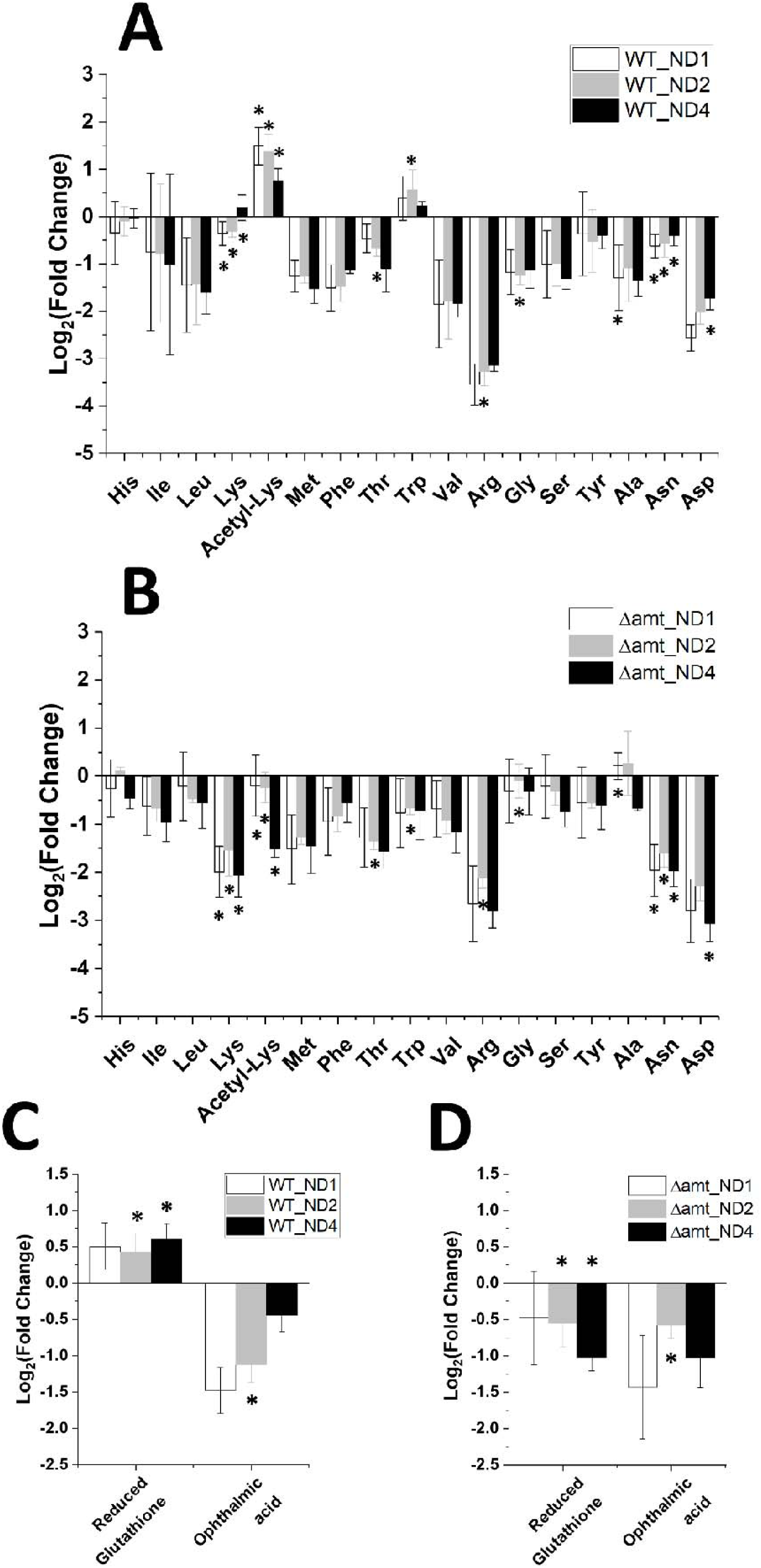
Pool of free amino acids and oxidative stress markers in both WT and Δamt strain, upon N deprivation. Free amino acid pool (A and B) and oxidative markers (C and D) in the WT strain (A and C) and in the Δamt mutant (B and D) in the four time points chosen in this experiment, according to Fig 4A. Data are expressed as base two logarithm of the fold change (FC) of the abundance of each metabolite between each of the three time points after N deprivation (i.e. ND1, 2 and 4) and N replete conditions (NR). Data are indicated as average ± SD of 6 biological replicates. Statistically significant differences between WT and Δamt for each metabolite at a specific time point are indicated with an asterisk (one-way ANOVA, p-value < 0.05).

Overall, both strains display a reduction in the whole pool of free amino acids as a consequence of N deprivation (Fig. 9A and B), likely suggesting a faster turnover upon N-fixing conditions. Nevertheless, upon N deprivation, the Δ*amt* strain also shows substantial remodelling of the free amino acid pools (Fig. 9B), relative to the parental strain (Fig. 9A). Major amino acids affected by the mutation are Lys and Asn, which display a stronger reduction in the mutant upon N deprivation, followed by Thr, Trp, Arg, Gly, Ala and Asp (Fig. 9B), which instead show minor but still relevant alterations. These results indicate a comprehensive impact on the amino acid metabolism in 7120 as a consequence of the mutation. It is also worth noting the pool of acetyl-lysine is differentially regulated in the two strains upon N deprivation (Fig. 9A and B), suggesting a comprehensively different regulation of the whole central metabolism.

Amino acids are substrates for the synthesis of several molecular players, important for the homeostatic control of the cell. Among them, glutathione (Cameron & Pakrasi, 2010) and ophthalmic acid (Ito *et al*, 2018) belong to a robust antioxidant buffering system which plays an important role in protecting against reactive oxygen species (ROS) generated as by-product of photosynthetic metabolism (Narainsamy *et al*, 2016).

Interestingly, while no major differences between the two strains were observed for ophthalmic acid upon N deprivation, the content of reduced glutathione (GSH) increased in WT (Fig. 9C) and it decreased in Δ*amt* (Fig. 9D), suggesting the mutant suffers from redox stress upon N deprivation.

### Gln, Glu and 2-OG pools

The metabolic pool concentration of several key metabolites in N metabolism, Gln, Glu and 2-OG [Fig. 1, (Martín-Figueroa *et al*, 2000; Böhme, 1998; Picossi *et al*, 2005)], was also affected by the *amt* mutation. The pool of free Glu increased in both strains over the course of the experiment (Fig. 10A and B), whilst the concentration of Gln dropped substantially in the wild-type upon N deprivation (Fig. 10A). Interestingly, the concentration of Gln is several-fold lower in Δ*amt* under N replete conditions and also drops upon elimination of assimilable N (Fig. 10B). Hence, the Gln/Glu ratio in the mutant is indicative of a partially deprived N metabolic state even in the presence of assimilable N (Supplementary Fig. S7), potentially affecting also the metabolic exchange between the two cell types (no difference in the number of heterocysts was observed between the two strains over the course of the experiment, see Supplementary Fig. S4). Similarly, the mutant also has a slightly lower 2-OG content than the parental strain under NR conditions (Fig. 10C), and consequently a higher 2-OG/Gln ratio (Supplementary Fig. S8), in line with the hypothesis that 2-OG is indicative of metabolic N availability (Muro-Pastor *et al*, 2001). The difference between the two strains is admittedly small, at only around 18% – but then, the decrease in 2OG following N deprivation is only about 40%, so even this small decrease could represent a change in N status.

**Figure 10.**
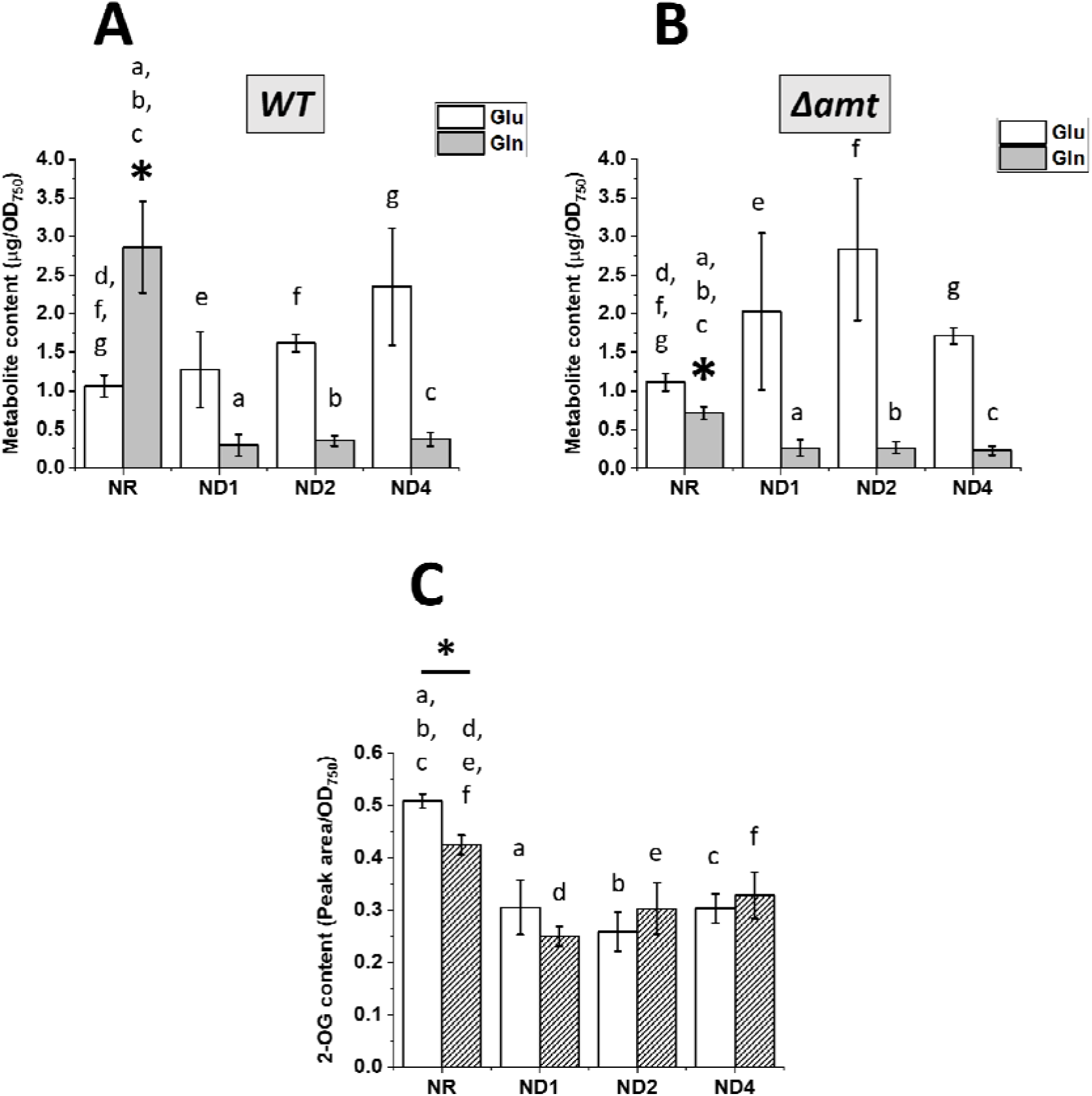
Metabolic pool concentration of key metabolites in N metabolism. Glu, Gln (A, B) and 2-OG (C) content in 7120 WT and Δamt strains, in the experimental conditions of Fig. 4A. Glu (white bars) and Gln (grey bars) content is split in two distinct panels for WT (A) and Δamt (B). C. 2-OG content in WT (white bars) and Δamt (striped bars). Results come from the same amount of biomass for both strains and for different growth conditions. Data are indicated as average ± SD of 6 biological replicates. Statistically significant differences between WT and Δamt for each metabolite at a specific time point are indicated with an asterisk, whilst the same alphabet letter indicates statistically significant differences for the same strain and metabolite, in different growth conditions (one-way ANOVA, p-value < 0.05).

## Discussion

Biological systems can be fuelled by multiple N sources, but they are all converted to ammonium/ammonia before assimilation, as the latter is the most reduced and energetically favourable bioavailable form of N. The translocation of ammonium/ammonia across biological membranes is therefore expected to play a potential key role in the regulation of N metabolism. Active ammonium/ammonia translocation [whether it involves the charged or uncharged form is still debated and it is likely to depend on the species in question (Ludewig, 2006; Ludewig *et al*, 2007; Boogerd *et al*, 2011; Wang *et al*, 2012; Javelle *et al*, 2007)] across biological membranes is catalysed by AMT transporters, a protein family widely distributed across multiple domains of life (Andrade & Einsle, 2007).

In this work we exploited a KO mutant of the whole *amt* cluster (i.e. *Δamt*) in *Anabaena sp. PCC 7120* (Paz-Yepes *et al*, 2008) to perturb both sensing and uptake of NH4^+^, with the aim to investigate how a N_2_-fixing cyanobacterium responds to perturbation of N -metabolism at a whole cell level.

We cultivated both 7120 WT and Δ*amt* strains in different N regimes (i.e. different N sources and N replete/deplete conditions). The underlying idea was to trigger different inner N states and investigate them through physiological, proteomic and metabolomic analyses. Upon N deprivation, *Anabaena sp. PCC 7120* differentiates a fraction of its cells into heterocysts to enable efficient N fixation (Golden & Yoon, 1998; Kumar *et al*, 2010), posing several limitations to integrated studies such as this one. One of them is the need to process the samples as a mixture of the two cell types, in order to avoid metabolic changes that are inevitable consequence of physical separation (Ermakova *et al*, 2014). This necessary choice forgoes discrimination of the metabolic status of the two cell types. Nevertheless, some proteins and metabolites are unique to one of the two cell types (Martín-Figueroa *et al*, 2000), enabling to partially overcome such limitations. Moreover, we did not observe any difference in the ratio of heterocysts to vegetative cells in response to any of the genetic or environmental treatments investigated in this work (Supplementary Fig. S4).

### Lack of AMT transporters triggers a substantial response at the whole cell level, but doesn’t induce any visible phenotype

In the constant laboratory conditions tested in this work, we observed AMT transporters are not essential to support growth of 7120, regardless of the N source used to sustain the central metabolism (Fig. 2A and 4A), as previously reported (Paz-Yepes *et al*, 2008). This finding corroborates what has been observed in other bacteria [e.g. *Rhodobacter capsulatus* (Yakunin & Hallenbeck, 2002)], suggesting a conserved transporter-independent function for AMT transporters across unrelated bacteria species.

Nevertheless, the mutant surprisingly displays several changes in the abundance of proteins and metabolite pools with a central role in N metabolism, triggering also a substantial effect on key enzymatic activities. Among them, we observed: (1) a substantial increase in nitrogenase activity (Fig. 2B and 5A), likely due to an increased accumulation of the protein complex (Figure 5B and 5C); (2) an increased GS activity in NH4^+^-replete conditions and upon prolonged N deprivation (Fig. 2C and 6A), as a consequence of changes in the abundance of proteins which play both a direct and/or indirect role on GS activity such as: (a) changes in the abundance of both the GS protein itself (Fig. 3A and 6B) and the post-translational negative regulator IF7A (Supplementary Fig. S3 and Fig. 6C) and (b) changes in the abundance of GDH, GOGAT and IDH, which re-generate its substrates (Supplementary Fig. S5). It is worth noting that some of these observations correlate with what has already been observed in other organisms, such as the photosynthetic purple bacterium *Rhodobacter capsulatus*, in which the absence of the AMTB transporter influenced both nitrogenase and GS activity, thus strengthening the hypothesis that such proteins might share the same role in the regulation of N metabolism, even in distinct bacteria species (Yakunin & Hallenbeck, 2002). Moreover, we also observed that the pool of free amino acids (Fig. 9A and 9B), redox markers (Fig. 9C and 9D) and metabolites with a key role in N metabolism orchestration (i.e. Gln, Glu and 2-OG, Fig. 10) is affected by the mutation. The *Δamt* mutant also displays substantial changes in photosynthetic performances and pigment content with respect to the parental strain (Fig. 4B and Table 1).

It is worth noting that the biochemical changes observed in the mutant do not translate in any phenotypic difference with respect to the parental strain (i.e. growth is unaffected, Fig. 2A and Fig. 4A), highlighting the strong robustness of the biological system under investigation. The latter most likely depends on its ability to undergo this very substantial homeostatic adjustment at the whole cell level, as a consequence of both genetic (i.e. *Δamt*) and environmental treatments (i.e. different N regimes).

### Lack of AMT transporters induces metabolic adaptation spanning both C and N metabolism, with a potential impact on the metabolites exchange between heterocysts and vegetative cells

As in many biological systems, N and C metabolism is expected to be tightly coupled (Zhang *et al*, 2018) also in 7120, thereby maintaining a properly balanced C/N ratio even when exposed to external perturbations (Forchhammer & Selim, 2019). When the C source (i.e. CO_2_ in case of phototrophic metabolism) is not limiting, external N source(s) are expected to directly influence the C/N balance of the cell, with both: (1) efficiency in sensing and uptake and (2) abundance and nature of such N source(s) playing a central role. In our experiments, 7120 was cultivated in a 1% CO_2_-enriched atmosphere in order to avoid C limitation and both genetic (i.e. *Δamt* mutation) and environmental (i.e. different N sources and abundance) treatments were used to perturb both such parameters and investigate the response of 7120 at a whole cell level.

The mutant displayed substantial changes in the abundance of several metabolites (Fig. 9), including Gln, Glu and 2-OG. Among them, it is worth noting the difference in the free pool of acetyl-lysine (Fig. 9A and 9B) which suggests the overall central metabolism regulation might be comprehensively affected as a consequence of the mutation (Nakayasu *et al*, 2017; Christensen *et al*, 2019). The observed differences in the free pool of acetyl-lysine might reflect either changes in total protein acetylation, or else changes in the turnover rates of acetylated proteins, with a potential regulatory role in both photosynthesis and carbon metabolism, as suggested in *Synechocystis sp. PCC 6803* (Mo *et al*, 2015). Further confirmation in the lab is however needed to clarify the regulatory role of protein acetylation in 7120.

Differences in the accumulation of Gln, Glu and 2-OG are a clear hallmark of a perturbed C/N balance (Fig. 10). Gln, Glu and 2-OG are in fact key metabolites involved in the GS-GOGAT cycle, hence they play a critical role in the central crossroad for C and N metabolism. Moreover, Gln and Glu metabolism is intertwined with that of several other amino acids as they are the most important amino groups donor for their synthesis (Reitzer, 2003; Huergo & Dixon, 2015), whilst 2-OG is the major signalling metabolite used to perceived the intracellular N status by cyanobacteria (Muro-Pastor *et al*, 2001). In our experiments, 2-OG concentration decreased upon N deprivation (Fig. 10C) and this correlates with the observed increase in the pool of free Glu (Fig. 10A and 10B), as cyanobacteria lack 2-OG dehydrogenases and therefore 2-OG is mainly used for the biosynthesis of Glu or other Glu-derived compounds (Herrero *et al*, 2001). It is worth noting that this trend is not affected by the mutation (Fig. 10C), which instead induces a reduction in the content of 2-OG in NH_4_^+^-replete conditions (Fig. 10C).

Taken together, these data suggest the absence of AMT transporters results in a metabolic adjustment in response to environmental treatments (i.e. different N regimes), and although not investigated in the present study, this is likely to affect also the traffic of metabolites between heterocysts and vegetative cells, as also suggested by the observed differences in CPG metabolism (Fig. 8).

### Are AMT transporters an integral part of the N metabolism regulatory/signalling network?

In our experiments, we repeatedly observed substantial changes in GS activity as a consequence of the mutation. The mutant in fact displays an increased GS activity in NH_4_^+^ replete and also in prolonged N deplete conditions (Fig. 6A and Fig. 2C), likely in response to changes in the abundance of both the GS protein itself and its post-translational regulator IF7A (Fig. 3 and 6). Moreover, we observed that GS activity is often retained even if IF7A abundance varies, potentially suggesting an additional molecular player(s) might also be involved in its regulation. Taken together, our results suggest AMT transporters might play a direct or indirect IF7A-independent role on GS activity in 7120, thus calling for further scientific efforts in order to fill potential gaps in the regulatory/signalling network of N metabolism.

The changes observed in this work are widespread at a whole cell level, also affecting the master molecular players which orchestrate N metabolism (i.e. NtcA, PII and PipX, Fig. 7). In particular, the mutant displays an increased PII abundance in NR conditions, with respect to the parental strain (Fig. 7B). PII belongs to one of the most widely distributed families of signal transduction proteins in nature, involved in various aspects of N metabolism and regulation of C/N homeostasis (Forchhammer, 2004, 2008; Arcondeguy *et al*, 2001; Forcada-Nadal *et al*, 2018). Among them, in heterotrophic bacteria and in archaea, PII proteins of the subfamily GlnK directly interact with AMT transporters to regulate their activity, typically reducing their uptake rate in N excess conditions to prevent intracellular over-accumulation of ammonium (Arcondeguy *et al*, 2001). Recent studies suggest PII protein binds AMT transporters also in cyanobacteria [i.e. *Synechococcus sp. PCC 7942* (Forchhammer & De Marsac, 1995) and *Synechocystis sp. PCC 6803* (Watzer *et al*, 2019)]. Our observation of a greater PII abundance, as a consequence of a deletion in AMT transporters, supports the hypothesis that PII might retain this function also in 7120. If true, this suggests one potential mechanism for how AMT transporters might influence the regulatory/signalling network of N metabolism in 7120.

It is worth noting that the molecular fingerprint of *Δamt* cells display symptoms of N - deficiency relative to the parental strain, also in NH_4_^+^ replete conditions, and that observations at both protein and metabolite level support this hypothesis. Among them: (1) the faster activation of nitrogenase upon N deprivation (Fig. 5A), (2) the increased GS activity (Fig. 6A), (3) the strong reduction in the Gln/Glu ratio (Supplementary Fig S7), (4) the slightly reduced 2-OG content (Fig. 10C) and (5) the increased 2-OG/Gln ratio (Supplementary Fig. S8). In contrast, the greater photosynthetic activity (Fig. 4B) and Chl/Car ratio (Table 1) suggest the mutant does not suffer from N limitation in NR conditions. This raises the interesting question: what is cause and what is effect? The observed changes at the protein level are coupled to a widespread metabolic adaptation as a consequence of the mutation, involving substrates and products of most of the enzymatic reactions investigated at the protein level. In fact, it is reasonable to expect that some of the changes in the accumulation of specific proteins might simply result from an adaptation to perturbations of the metabolite pool or *vice versa*. At this point, we thus cannot conclude whether the *Δamt* mutant either is directly suffering from an imbalanced regulatory/signalling network or if it suffers from perturbations at the metabolite level, which are sensed by such regulatory/signalling network. However, AMT transporters play a direct or indirect role in the regulatory/signalling network which still warrant further investigations.

## Conclusions

We investigated how a mutant of *Anabaena sp. PCC 7120*, impaired in both sensing and low-affinity up-take of NH_4_^+^, responds to environmental treatments affecting the inner N status of the cells. The whole cell system responds with substantial internal perturbations embracing both N and C metabolism. Moreover, the absence of AMT transporters leaves a molecular fingerprint suggesting N-deficiency, which surprisingly does not lead to any externally measurable phenotypic effects. We thus hypothesise that 7120 evolved a robust regulatory/signalling molecular network to maintain N metabolism homeostasis. The observed changes involve both proteins and metabolites, highlighting a pleiotropic effect of the mutation. We also provided evidence of perturbations to nitrogenase and GS activity, as well as to master regulators orchestrating N metabolism, thus leading to the hypothesis that a possible direct or indirect IF7A-independent role of AMT transporters on GS activity exists in 7120, possibly transduced via the PII protein. Taken together, these evidences suggest AMT transporters might play an active role in the regulatory/signalling network, calling for further scientific efforts in order to fill current gaps in N metabolism homeostasis. The work highlights the dynamic and complex nature of internal mechanisms involved in maintaining homeostasis and the success in so doing, achieving a near-complete lack of any measurable external impact.

## Material and Methods

### Cyanobacteria strains and growth conditions

Strains of *Anabaena sp. PCC 7120* used in this work are summarised in table 2 and were kindly provided by Prof. Enrique Flores [institute of Plant Biochemistry and Photosynthesis, University of Sevilla (Sevilla, Spain)].

**Table 2.** Strains of Anabaena sp. PCC 7120 used in this work.

| Strain | Genotype | Source |
| --- | --- | --- |
| *Anabaena sp. PCC 7120* WT | Wild-type (WT) | Prof. Enrique Flores |
| *Anabaena sp. PCC 7120 Δamt* | Δamt::C.K3;<br>Amt4<sup>-</sup>, Amt1<sup>-</sup>, AmtB<sup>-</sup>, Nm<sup>R</sup> | (Paz-Yepes *et al*, 2008) |

Both strains were maintained in solid BG11 medium (pH 8.0 with the addition of 10 mM final concentration of TES-NaOH buffer) (Rippka *et al*, 1979) (1.5% Difco Bacto agar, BD) in an Algaetron 230 (PSI, Photon Systems Instruments, Czech Republic), with an atmosphere enriched in CO_2_ (1%), under cool white light at 60 µmoles of photons*m^-2^*s^-1^, 30 °C. Illumination rate was determined using a LI-250A photometer (Heinz-Walz, Effeltrich, Germany). Before starting an experiment, both strains were switched to liquid medium and pre-cultivated axenically in 100-ml Erlenmeyer flasks in 20 ml BG11_0_ (BG11 medium, without NaNO_3_) supplemented with 5 mM NH_4_Cl in the same growth conditions, under orbital shaking at 160 rpm. Cells were kept in exponential growth conditions, refreshing the cultures every other day (i.e. replacing half of the volume of the culture with fresh BG11_0_ + 5 mM NH_4_Cl). *Anabaena sp. PCC 7120* Δ*amt* strain was cultivated in presence of 10 µg/ml neomycin (Nm).

For all experiments carried out in this work, strains were cultivated in 6-well polystyrene plates in the same growth conditions indicated above. Cells pre-cultivated in Erlenmeyer flasks were washed twice in the final growth medium, through centrifugation for 10 min, 3500 g, RT. Starting OD_750_ for all growth curves = 0.1. Media at different pH values were obtained using 10 mM final concentration of phosphate, TES-NaOH and CAPS buffers, respectively for pH 6.0, 8.0 and 10.0.

Growth was monitored through OD_750_ in 96-wells polystyrene plates with a multimode spectrophotometer (Tecan Infinite M200 Pro). Linear correlation between OD_750_ and biomass dry weight was confirmed for both strains and the growth ranges measured in this work. Biomass dry weight was measured gravimetrically as previously reported in (Perin *et al*, 2015). Specific growth rate was calculated by the slope of different growth phases for growth curves plotted in logarithmic scale.

### Pigments content and photosynthetic efficiency

Pigments from intact cells grown in 6-wells plates were extracted using a 1:1 biomass to solvent ratio of 100% methanol, at 4 °C in the dark for at least 20 min (Sinetova *et al*, 2012). Absorbance at 470, 665 and 730 nm was monitored using a multimode spectrophotometer (Tecan Infinite M200 Pro) to determine pigment concentrations, using specific extinction coefficients (Ritchie, 2006; Wellburn, 1994).

Photosynthetic efficiency was assessed measuring *in vivo* chlorophyll fluorescence of intact cells using an AquaPen-C AP 110-C (PSI, Photon Systems Instruments, Czech Republic). Photosystem II (PSII) functionality was assessed as PSII maximum quantum yield (Φ_PSII_), according to (Maxwell & Johnson, 2000).

### Ammonium/ammonia quantification

Ammonium/ammonia quantification was performed using an adaptation of the method described by Willis *et al*. (Willis *et al*, 1996) in order to enable the use of 96-well microtiter plates. Briefly, *Anabaena sp. PCC 7120* strains grown in 6-well plates were harvested by centrifugation and 10 µl of the cell-free supernatant were loaded in a flat-bottom 96-well plate. 200 µl of reactive solution (32 g/L sodium salicylate, 40 g/L Na_3_PO_4_·12H_2_O and 0.5 g/L sodium nitroprusside) and 50 µl of hypochlorite solution (0.25-0.37% active chlorine) were added in this order to the cell-free sample and the solution made homogenous, before measuring the absorbance at 685 nm in a multimode spectrophotometer (Tecan Infinite M200 Pro), after incubation for 15 min, 900 rpm, RT in a shaker (PHMP, Grant Instruments). Ammonium/ammonia concentration in the cell-free samples was calculated using the linear range of a standard curve prepared with serial dilutions of a NH_4_Cl solution.

### Enzymatic activities

#### Glutamine synthetase

Glutamine synthetase (GS) activity was assessed through the method detailed below, which comes from the combination of different protocols from (Bressler & Ahmed, 1984; Orrs *et al*, 1981; Merida *et al*, 1991).

##### Preparation of cell-free total proteins extracts

Strains of *Anabaena sp. PCC 7120* grown in 6-wells plates were harvested by centrifugation and the supernatant discarded. Cell pellets were transferred to 2 ml polypropylene tubes and disrupted twice using a TissueLyser II (Qiagen), adding the same volume of acid-washed glass beads (Sigma-Aldrich) and solubilisation buffer (50 mM Hepes, 0.2 mM EDTA, pH 7.3), for 5 min at 30 Hz. Tube holders were pre-cooled at -20 °C. Cell extracts were collected through centrifugation for 10 s, 9000 g, 4 °C. Cell extracts were centrifuged twice for 10 min, 21000 g, 4 °C to eliminate cell debris and insoluble proteins. Total proteins concentration in the cell-free lysate was assessed through DC protein assay (BIO RAD), following the manufacturer’s manual, using 96-wells plates against a BSA (bovine serum albumin) standard curve. *GS activity assay*. GS catalyses the condensation of glutamate and ammonia to generate glutamine, hydrolysing ATP (reaction below).

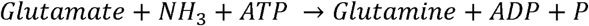

GS activity can be measured indirectly, quantifying the amount of phosphate released by the reaction through colorimetric assay.

Enzymatic assay was performed in 550 µl in 24-wells flat bottom polystyrene plates, with 50 µg total proteins in 364 mM imidazole-HCl, pH 7.0, 1.82 mM NH_4_Cl, 5.45 mM Na-ATP·H_2_O, pH 7.0 (prepared fresh on ice), 0.52 M MgCl_2_·6H_2_O, 91 mM sodium glutamate, pH 7.0, for 15 min, 400 rpm, 30 °C in a thermoshaker (PHMP, Grant Instruments). The reaction was then quenched on ice, adding FeSO_4_·6H_2_O (0.61% w/v in 0.011 N H_2_SO_4_ final concentration). In order to develop colour (proportional to the amount of phosphate released by ATP as a consequence of GS activity), ammonium heptamolybdate (0.4% w/v in 0.45 M H_2_SO_4_ final concentration) was added to the reaction and solutions were homogenised on ice. Colour intensity of clear samples was measured at 850 nm, using a multimode spectrophotometer (Tecan Infinite M200 Pro). GS activity was calculated subtracting to each sample the OD_850_ of the corresponding blank solution (prepared replacing the substrate sodium glutamate with water and corresponding to the background signal of phosphate released by other endogenous ATP-dependent enzymatic reactions).

#### Nitrogenase

Nitrogenase activity was determined with an acetylene reduction assay under oxic culture conditions. Cells grown in 6-well plates were incubated in 2-ml gas chromatography glass vials under an atmosphere of 10% acetylene in air. Vials were incubated for 3 h in the same original growth conditions (i.e. shaking, light, 30 °C) and the quantity of ethylene in the headspace was determined by gas chromatography (7820A GC system, Agilent Technologies). The nitrogenase activity is expressed as % conversion of added acetylene into ethylene, per hour and per µg of Chl.

### Cyanophycin content

#### Cyanophycin extraction

The same amount of biomass (OD_750_ = 0.3) of different *Anabaena sp. PCC 7120* strains grown in 6-wells plates was harvested by centrifugation and the supernatant discarded. Cyanophycin was extracted from the cell pellets following the protocol detailed in (Watzer *et al*, 2015), with some modifications as follows. Briefly, cell pellet was resuspended in 1 ml 100% acetone and incubated in a shaker for 30 min, 1400 rpm, RT. Lysed cells were then centrifuged for 17 min, 21000 g, RT and the supernatant discarded. Pellet was resuspended in 1.5 ml of 0.1 M HCl and incubated for 1 h, 1400 rpm, 60 °C to solubilise cyanophycin polymers. Solubilised cyanophycin was centrifuged for 17 min, 21000 g, RT to remove immiscible debris. Tris-HCl, pH 8.0 was added to the supernatant (0.2 M final concentration) and samples incubated for 40 min, 4 °C. Samples were then centrifuged for 17 min, 21000 g, 4 °C and pelleted cyanophycin polymers were resuspended in 500 µl 0.01 M HCl for quantification.

#### Cyanophycin quantification

Cyanophycin is a polymer of arginine and aspartate (Forchhammer & Watzer, 2016) and the quantification of arginine released by cyanophycin granules can be used as proxy for cyanophycin content determination (Burnat *et al*, 2014). Arginine quantification was performed through a modified colorimetric Sakaguchi method, according to (Messineo, 1966).

### Proteomic analysis

#### Sample preparation

Strains of *Anabaena sp. PCC 7120* grown in 6-wells plates were harvested by centrifugation and the supernatant discarded. Cell pellets were transferred to 2 ml polypropylene tubes and disrupted using a TissueLyser II (Qiagen), adding the same volume of acid-washed glass beads (Sigma-Aldrich) and extraction buffer (20 mM Tris-HCl, pH 8.0, 1 mM EDTA, pH 8.0 and 2 mM DTT), for 5 min at 30 Hz. Tube holders were pre-cooled at -20 °C. Cells disruption was repeated twice and cell extracts were collected through centrifugation for 10 s, 9000 g, 4 °C and pulled together after both cycles. Cell extracts were centrifuged twice for 10 min, 21000 g, 4 °C to eliminate cell debris and insoluble proteins. Total proteins concentration was assessed through DC protein assay (BIO RAD), following the manufacturer’s manual, using 96-wells plates against a BSA (bovine serum albumin) standard curve. Cell extracts were diluted by 0.1 M ammonium bicarbonate to have a 0.3 µg/µl total proteins final concentration and the standard peptide [Glu1]-Fibrinopeptide B, human (GluFib, Sigma Aldrich) was added (0.3 ng/µl, final concentration). Samples were reduced by DTT and ammonium bicarbonate (final concentration 10 mM and 50 mM, respectively) for 1 h, 56 °C, and subsequently alkylated adding iodoacetamide prepared fresh in 0.1 M ammonium bicarbonate (50 mM final concentration), for 30 min, 37 °C, 500 rpm in a thermoshaker (PHMT; Grant Instruments). Alkylated samples were digested by proteomics-grade trypsin (Promega), final concentration 6 ng/µl, ON, 37 °C. Tryptic digestion was stopped quenching the reaction with formic acid (final concentration 1%) for 30 min, 37 °C, 500 rpm in a thermoshaker. Acidified samples were centrifuged at 17000 g, RT, to pellet water immiscible degradation products and the supernatant collected for mass spectrometry analysis.

#### Mass spectrometry

Trypsin digested samples were analysed on an AB-SCIEX 6500QTrap MS coupled to an Agilent 1100 LC system. Chromatography was performed on a Phenomenex Luna C18(2) column (100 mm x 2 mm x 3 µm) at 50 °C, using a gradient system of solvents A (94.9% H_2_O, 5% CH_3_CN and 0.1% formic acid) and B (94.9% CH_3_CN, 5% H_2_O and 0.1% formic acid). A gradient from 0 to 35% B over 30 min at a flow rate of 250 ml/min was used. The column was then washed with 100% B for 3 min and then re-equilibrated with 100% A for 6 min. Typically 40 µl injections were used for the analysis. The MS was configured with an Ion Drive Turbo V source; Gas 1 and 2 were set to 40 and 60, respectively; the source temperature to 500 °C and the ion spray voltage to 5500 V. MS, configured with high mass enabled, was used in “Trap” mode to acquire Enhanced Product Ion (EPI) scans for peptide sequencing and in “Triple Quadrupole” mode for Multiple Reaction Monitoring (MRM). Data acquisition and analysis was performed with SCIEX software Analyst 1.6.1 and MultiQuant 3.0. Signature peptides for all the proteins investigated in this work were determined through trial MRM runs. Only the peptide GluFib was used as standard for protein normalisation. The typical work-flow to select the best signature peptides involved the analysis of trial samples in different growth conditions using transitions generated by *in silico* analysis with the open-source Skyline Targeted Mass Spec Environment (MacCoss Lab) (MacLean *et al*, 2010; Abbatiello *et al*, 2013). The identity of candidate peptides was then confirmed by EPI scans. Background proteome of *Anabaena sp. PCC 7120* (http://genome.kazusa.or.jp/cyanobase) was used to check for uniqueness of target peptides. Typically, 3-5 transitions per peptide were used. The final method includes 1-4 peptides per protein for unique identification and quantification. Signature peptides for all the proteins investigated in this work are listed in table 3.

**Table 3.**
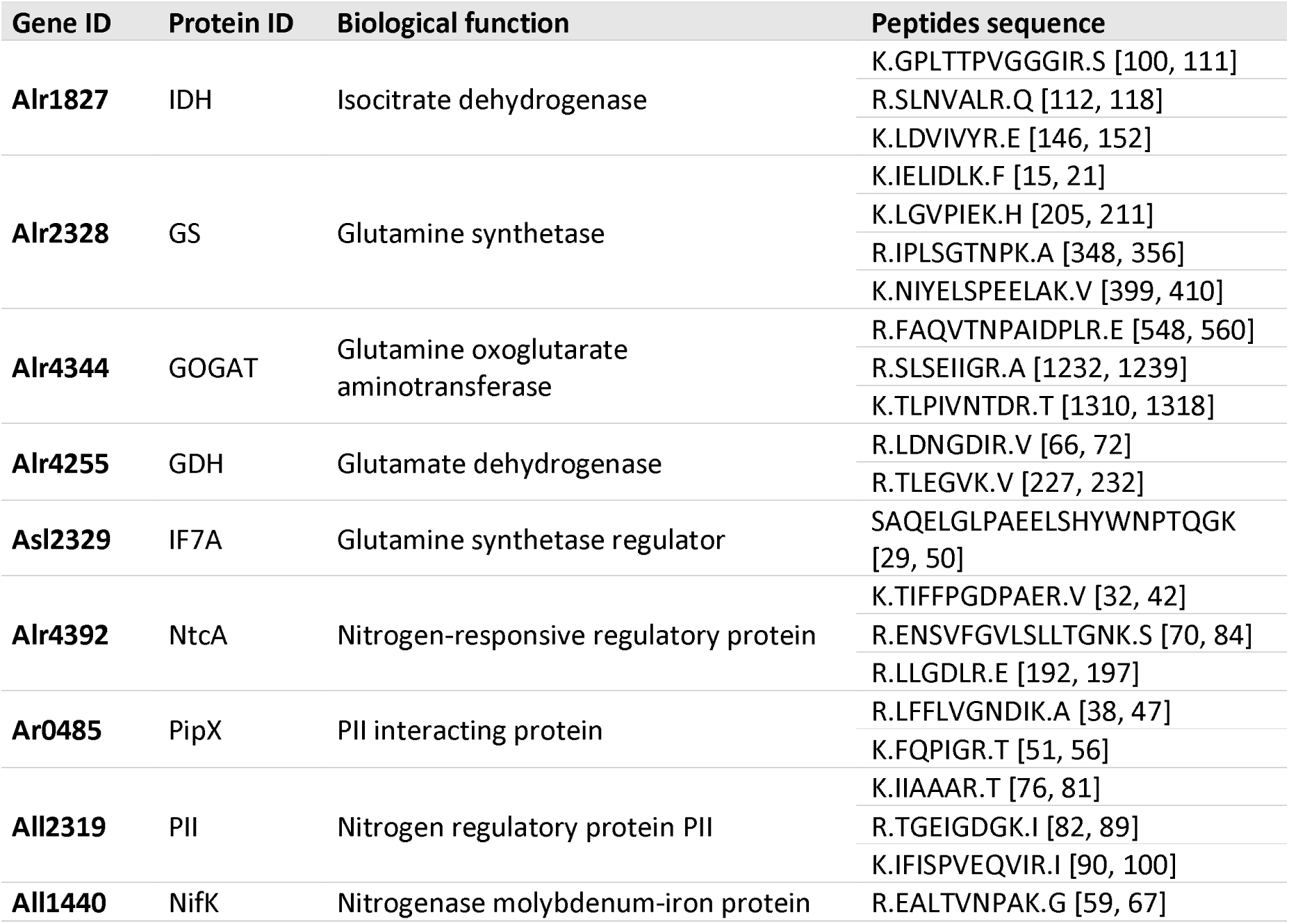
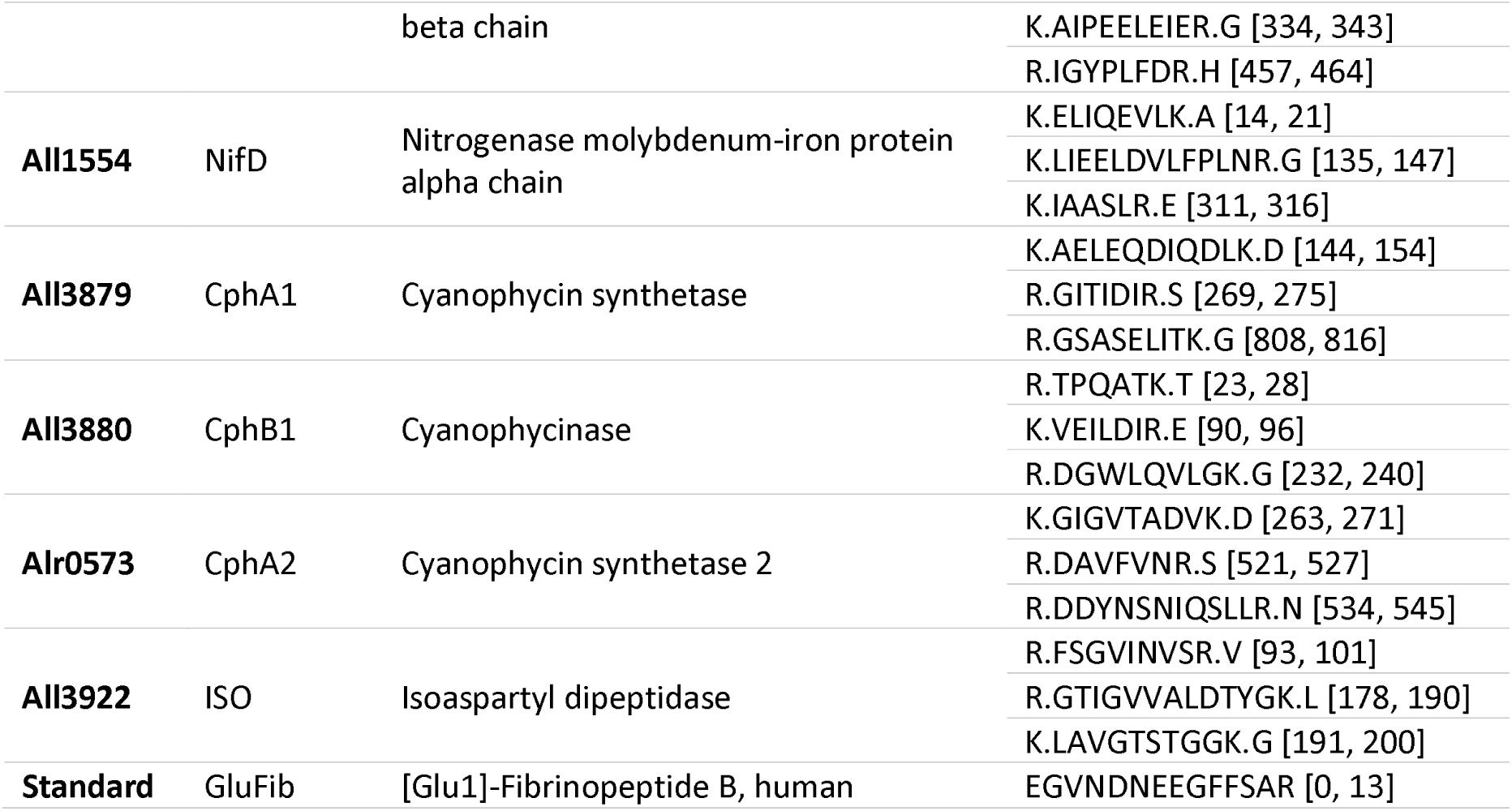
Signature peptides for the proteins investigated in this work. Gene ID, protein ID and corresponding biological function are indicated. Peptides sequences show their position within the corresponding protein in square brackets.

Protein quantification was performed accounting for the intensities of all transitions peaks for all the peptides belonging to a specific protein. The resulted peak area was normalized to the peak intensity of the GluFib peptide standard.

### Metabolomic analysis

#### Sample preparation

Strains of *Anabaena sp. PCC 7120* grown in 6-wells plates were harvested by fast filtration, modifying the protocol from (Eisenhut *et al*, 2008). Briefly, cells were fast filtered in the light without any subsequent washing step, using a vacuum filtration system (0.45 µm pore size nitrocellulose filter, 47 mm diameter (Sigma Aldrich)), using stainless-steel stand and funnel (Sartorius). Filters were then transferred to 50 ml tubes and immediately frozen in liquid nitrogen and stored at -80 °C until metabolites extraction. Time between harvesting and metabolic inactivation by freezing was <10 sec. Deep frozen cells were scraped off the nitrocellulose filters using 80% cold methanol (−20 °C). Cells in cold methanol were transferred to 2 ml polypropylene tubes and metabolite extraction was carried out with a TissueLyser II (Qiagen), using tube holders pre-cooled at -20 °C and adding the same volume of acid-washed glass beads (Sigma-Aldrich), for 5 min at 30 Hz. Metabolites were collected after centrifugation for 10 min, 21000 g, 4 °C. Metabolite extraction was repeated twice, the extracts pooled together and centrifuged again to separate cell debris and other immiscible products. Metabolic extracts were then dried by vacuum centrifugation overnight and stored at -80 °C until use.

#### Sample derivatisation and mass spectrometry

Dried metabolic extracts were reconstituted in 300 µl of water, using L-phenylalanine-d_5_ as internal standard (final concentration 2 μg/ml), then vortexed and centrifuged at 16000g for 10 min. Quality control (QC) samples were prepared by mixing 10 μl of the supernatant of each sample, with the analytical batch including 10% QC samples.

##### Amino acids quantification

6-aminoquinolyl-N-hydroxysuccinimidyl carbamate was used for derivatization (AccQTag derivatization) according to the manufacturer’s manual (AccQTag, Waters Corp). Briefly, 70 μl of borate buffer (pH 8.6) was added to 10 μl sample, followed by the addition of 20 µl AccQTag reagent (in acetonitrile). Samples were then vortexed and heated at 55°C for 10 minutes. 5 μl of each sample were analysed by HPLC-electrospray ionisation/MS-MS using a Shimadzu UFLC XR/AB SCIEX Triple Quad 5500 system, running in multiple reaction monitoring (MRM) via positive ionization mode. The LC-MS method here exploited is based on the one previously described by (Gray *et al*, 2017), with some modifications, as detailed below.

##### LC method

Mobile phase A: LC-MS grade water with 0.5% formic acid; mobile phase B: LC-MS grade acetonitrile with 0.5% formic acid. Flow rate: 300 μl/min, with the following gradient elution profile: 0.1 min, 4% B; 10 min, 28% B; 10.5 min, 80% B; 11.5 min, 80% B; 12 min, 4% B; 13 min,4% B. An Acquity HSS T3 UPLC column [2.1 mm × 100 mm, 1.8 μm particle size (Waters Corporation, Milford, MA, U.S.A.)] was used to achieve metabolite separation at 45°C.

##### MS method

The data were acquired through a Sciex QTRAP 5500 MS/MS system [Applied Biosystem (Forster City, CA, USA)], using the following settings: curtain gas, 40 psi; collision gas, medium; ionspray voltage, 5500 V; temperature, 550°C; ion source gas 1 and 2, 40 and 60 psi, respectively. De-clustering, entrance and collision cell exit potentials were set to 30 V, 10 V and 10 V, respectively. Multiple reaction monitoring (MRM) transitions, retention time (RT) and individually optimised collision energy (CE) for each metabolite are listed in supplementary table S1.

##### 2-oxoglutarate (2-OG) quantification

The ion pairing LC method here exploited was adapted from (Michopoulos, 2018), according to the following settings. Mobile phase A: 10 mM tributylamine (TBA) and 15 mM acetic acid in LC-MS grade water; mobile phase B: 80% methanol and 20% isopropanol. Flow rate: 400 μl/min, with the following gradient elution profile: 0 min, 0% B; 0.5 min, 0% B; 4 min, 5% B; 6 min, 5% B; 6.5 min, 20% B; 8.5 min, 20% B; 14 min, 55% B; 15 min, 100% B; 17 min, 100% B; 18 min, 0% B; 21 min 0% B. The method was transferred to XEVO TQ-S (Waters, Wilmslow, U.K.), using an Acquity UPLC system with 45 °C separation temperature. MS data were acquired according to the following parameters: capillary voltage, 0.8 kV; source offset, 50 V; desolvation temperature, 500°C; source temperature, 150°C; desolvation gas flow, 1000 L/h; cone gas flow, 150 L/h; neutraliser gas, 7.0 bar; collision gas, 0.15 ml/min; cone voltage, 80 V. Data were acquired through an electrospray negative ionisation, focusing only on [M-H]^-^ ions. Retention time (RT), multiple reaction monitoring (MRM) transitions and individually optimised collision energy (CE) are listed in supplementary table S2.

##### Data processing

The raw LC-MS data were analysed using Skyline [MacCoss Lab, (Adams *et al*, 2020)]. External dilution curves were used to determine the range for linear response.

##### Chemicals and reagents

Metabolite standards (Mass Spectrometry Metabolite Library of Standards, MSMLS) were purchased from IROA Technologies (Michigan, MI, U.S.A.). Acetic acid, tributylamine (TBA), L-phenyl-d5-alanine, were obtained from Sigma-Aldrich (Gillingham, U.K.). LC-MS grade water, water with 0.1% formic acid (v/v) and acetonitrile with 0.1% formic acid (v/v) were purchased from Fisher Scientific (Leicester, U.K.). Methanol and isopropanol were purchased from Honeywell (Charlotte, NC, U.S.A.). AccQTag Ultra reagent was purchased from Waters UK.

### Statistical analysis

Descriptive statistical analysis was applied for all the data presented in this work. Statistical significance was assessed by one-way analysis of variance (One-way ANOVA) using OriginPro 2018b (v. 9.55) (http://www.originlab.com/). Samples size was at least >4 for all the measurements collected in this work.

## Acknowledgments

We acknowledge professor Enrique Flores from the institute of Plant Biochemistry and Photosynthesis, University of Sevilla (Sevilla, Spain), for kindly providing both the *Anabaena sp. PCC 7120* wild-type (WT) and the KO strain for the *amt* gene cluster.

We also acknowledge help and support from Dr. Mark Bennett and Dr. Paul Hitchen with MRM-MS protein quantification, and from Dr. David Malatinszky for setting up high-throughput GS and ammonium quantification assays.

We thank Stephen Rothery (FILM) for help with image analysis also acknowledge the funding from the Wellcome Trust (grant 104931/Z/14/Z) and BBSRC (grant BB/L015129/1) which support the Facility for Imaging by Light Microscopy (FILM) at Imperial College London.

## Author’s contribution

GP and PJ, conception, design and writing of the manuscript. GP, experiments coordination, sample collection, data collection and analysis. TF, set up the method for MRM protein quantification and sample run. VSK, set up the method for metabolomic analysis and sample run. DG, set up ImageJ macro for microscope analysis. MC, sample preparation for metabolomic analysis. JB, metabolomic analysis coordination. All authors revised the manuscript and approved its final version.

