## Supplementary material for "Calm on the surface, dynamic on the inside. Molecular homeostasis in response to regulatory and metabolic perturbation of *Anabaena* sp. PCC 7120 nitrogen metabolism"

### Supplementary Materials and Methods

#### Microscopy analysis for heterocysts quantification

Filaments were visualised through LED illumination with a widefield Zeiss Axio Observer inverted microscope. Chl a was excited with a 488 nm argon ion laser, and fluorescence was collected across 680–720 nm. Heterocysts quantification is expressed as percentage of heterocysts in the filament and data for each biological replicate come from the analysis of 10 microscope fields of view. Image analysis was performed through the software ImageJ (v. 1.52; https://imagej.nih.gov/ij/index.html). In this work we developed an ImageJ macro to enable automatic detection of size ($\mu m^2$) and Chl a fluorescence (mV) for all the cells of the filaments in the microscope field (see Text 1). Only cells with the following parameters: size > 10 $\mu m^2$ and Chl a fluorescence < 3000 (mV) were counted as heterocysts (Supplementary Fig. S1).

***Text 1. ImageJ macro to enable automatic detection of size ($\mu m^2$) and Chl a fluorescence (mV) for all the cells of the filaments in the microscope field.***

```
////////////////////////////////////////////////////
//// Name: Cyanobacteria counter
//// Author: David Gaboriau and Stephen Rothery, Facility for Imaging by Light Microscopy (FILM),
Imperial College London
//// Version: 1.0
////
//// Usage: Segment cells in strings of Cyanobacteria imaged as z-stacks on the brightfield channel
and measure their size, and measure the intensity of the autofluorescence of their Chlorophyll
content.
////
/////////////////////////////////////////////////////



//select directory of images to open
input = getDirectory("Input directory where images are stored");

//select location where images/results are to be stored
output = getDirectory("Output ditrectory for results");

//gets list of files
list = getFileList(input);

//loop for opening images
for (image=0;image<list.length;image++){

full = input +  list[image];
//open(full);
run("Bio-Formats Importer", "open=full autoscale color_mode=Default view=Hyperstack
stack_order=XYCZT");

fn=getTitle();
```

```
run("Clear Results");

roiManager("reset");


//duplicate and project the BF stack
run("Duplicate...", "duplicate channels=2");
run("Z Project...", "projection=[Max Intensity]");
//invert and subtract background
run("Invert");
run("Subtract Background...", "rolling=5");


//first threshold to find edges and make binary
setAutoThreshold("Huang dark");
setOption("BlackBackground", false);
run("Convert to Mask");
// threshold inner parts and add to roi manager
setAutoThreshold("Default dark");
run("Analyze Particles...", "size=0-20 add");
resetThreshold();
//create a selection from rois
n=roiManager("count");
roiManager("Select", Array.getSequence(n));
roiManager("Combine");
roiManager("Add");
roiManager("delete");
//fill rois
roiManager("Select", 0);
setForegroundColor(0, 0, 0);
roiManager("Fill");

newname = replace(fn, ".czi", "");

//watershed and clean up image
run("Watershed");
run("Remove Outliers...", "radius=5 threshold=0 which=Dark");

roiManager("reset");
//second threshold to detect cells and add to roi manager
setAutoThreshold("Default");
run("Set Measurements...", "area mean min centroid perimeter feret's display add redirect=None
decimal=3");
run("Analyze Particles...", "size=100-850 pixel circularity=0.75-1.00 show=Outlines add");
saveAs("tif", output+newname+" Outlines.TIFF");
resetThreshold();
close();

selectWindow(fn);
run("Duplicate...", "duplicate channels=1");
run("Z Project...", "projection=[Max Intensity]");
```

```
roiManager("deselect");
roiManager("measure");
roiManager("Show All without labels");
saveAs("tif", output+newname+" Cells_Chlorophyl.TIFF");


selectWindow(fn);
run("Duplicate...", "duplicate channels=2");
run("Z Project...", "projection=[Max Intensity]");
roiManager("deselect");
roiManager("Show All without labels");
saveAs("tif", output+newname+" Cells_Brightfield.TIFF");



selectWindow("Results");
saveAs("Results", output+newname+" Results.xls");

run("Close All");



}

run("Close All");
```

**Supplementary tables**

***Supplementary Table S1. Multiple reaction monitoring (MRM) transitions, retention time (RT) and individually optimised collision energy (CE) for each metabolite applied in the amino acids quantification method with 6-aminoquinolyl-N-hydroxysuccinimidyl.***

| Compound | Parent (m/z) | Fragment(m/z) | RT (min) | CE (eV) |
|---|---|---|---|---|
| **Lysine** | 244.06 | 171.1 | 5.9 | 20 |
| **Glycine** | 246.03 | 171.1 | 3.3 | 30 |
| **Alanine** | 260.05 | 171.1 | 4.4 | 30 |
| **Serine** | 276.04 | 171.1 | 2.8 | 30 |
| **Valine** | 288.08 | 171.1 | 6.9 | 25 |
| **Threonine** | 290.06 | 171.1 | 4.1 | 30 |
| **Leucine** | 302.09 | 171.1 | 8.6 | 30 |
| **Isoleucine** | 302.09 | 171.1 | 8.4 | 30 |
| **Asparagine** | 303.05 | 171.1 | 2.6 | 30 |
| **Aspartate** | 304.04 | 171.1 | 3.6 | 30 |
| **Glutamine** | 317.07 | 171.1 | 3 | 30 |
| **Glutamate** | 318.05 | 171.1 | 3.4 | 30 |
| **Methionine** | 320.05 | 171.1 | 6.8 | 30 |
| **Histidine** | 326.07 | 171.1 | 1.8 | 30 |
| **Phenylalanine** | 336.1 | 171.1 | 8.9 | 30 |
| **Arginine** | 345.11 | 171.1 | 2.64 | 35 |
| **Tyrosine** | 352.07 | 171.1 | 6.6 | 30 |
| **Acetyl-lysine** | 359.12 | 171.1 | 4.9 | 40 |
| **Tryptophan** | 375.09 | 171.1 | 9.4 | 30 |
| **Reduced Glutathione** | 478 | 171.1 | 5.2 | 30 |
| **Ophthalmic acid** | 460 | 171.1 | 4.4 | 35 |

***Supplementary Table xx. 2-oxoglutarate (2-OG) quantification****. Multiple reaction monitoring (MRM) transitions, retention time (RT) and individually optimised collision energy (CE) for each metabolite applied in the ion-pairing LC method.*

| Compound | Parent (m/z) | Fragment(m/z) | Rt (min) | CE(eV) |
|---|---|---|---|---|
| **2-oxoglutarte (2-OG)** | 145.01 | 101.1 | 8.7 | 13 |
| | | 57 | 8.7 | 11 |
| | | 127 | 8.7 | 14 |
| **2-oxoglutarte (2-OG) labelled $^{13}C_4$** | 149.01 | 104.1 | 8.7 | 11 |

**Supplementary figures**

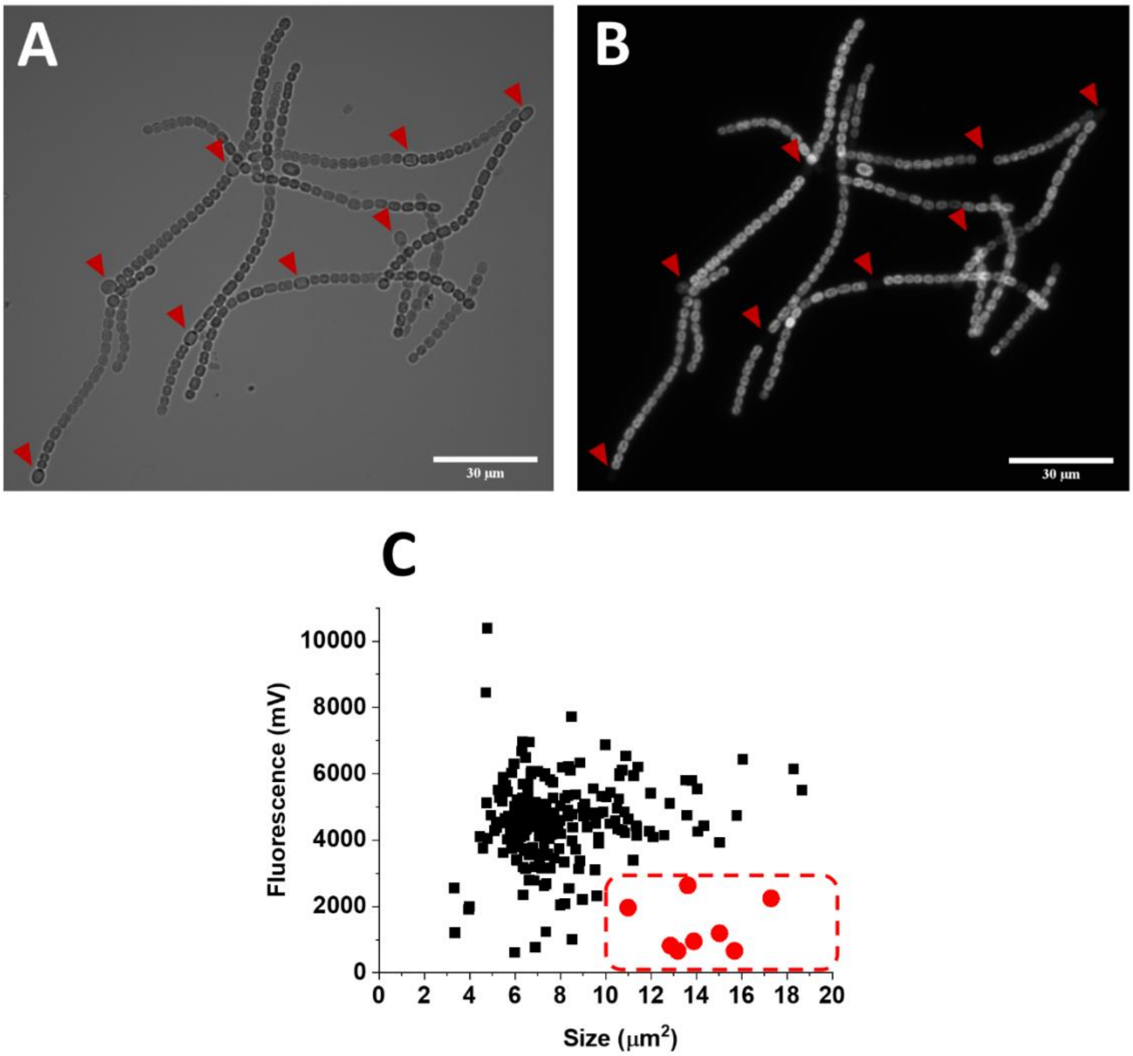


***Supplementary figure S1. Validation of the heterocysts counting protocol.*** *A and B show brightfield and Chl a autofluorescence microscope images, respectively, for Δamt filaments in ND4 (4 days after N deprivation). Red arrows indicate the heterocysts identified by the macro described in Text 1, which correspond to the red circles in C, where size of cells of the filaments in A and B is plotted against their corresponding Chl a fluorescence values. The red dashed square highlights the region of the plot meeting the heterocysts selection criteria as indicated in supplementary Materials and Methods.*

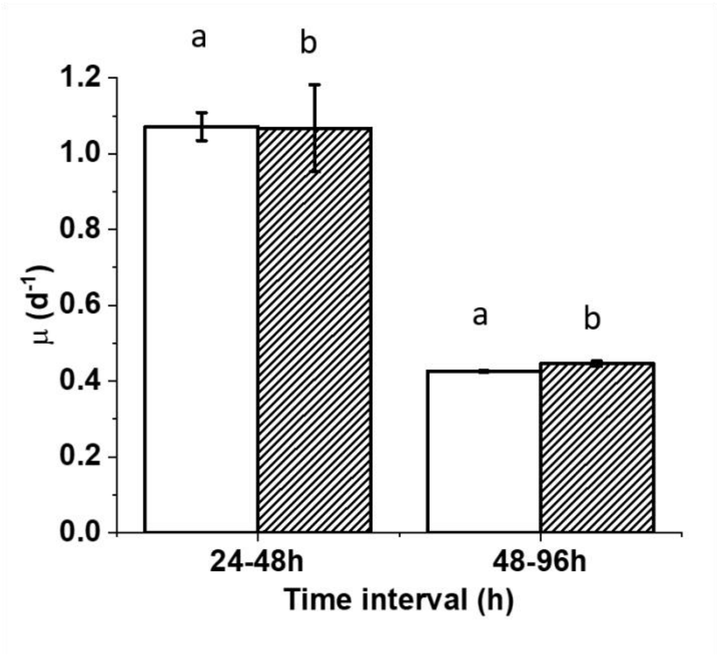


***Supplementary figure S2. Specific growth rate (μ) in different time intervals for 7120 WT and Δamt strains grown in $BG11_0$ + 5 mM $NH_4^+$.*** *Growth curves used to calculate this parameter are shown in figure 2A. Data are indicated as average ± SD of 6 biological replicates. Statistically significant differences between WT (white bars) and Δamt (striped bars) are indicated with an asterisk, whilst the same alphabet letter indicates statistically significant differences for the same strain in different growth conditions (one-way ANOVA, p-value < 0.05).*

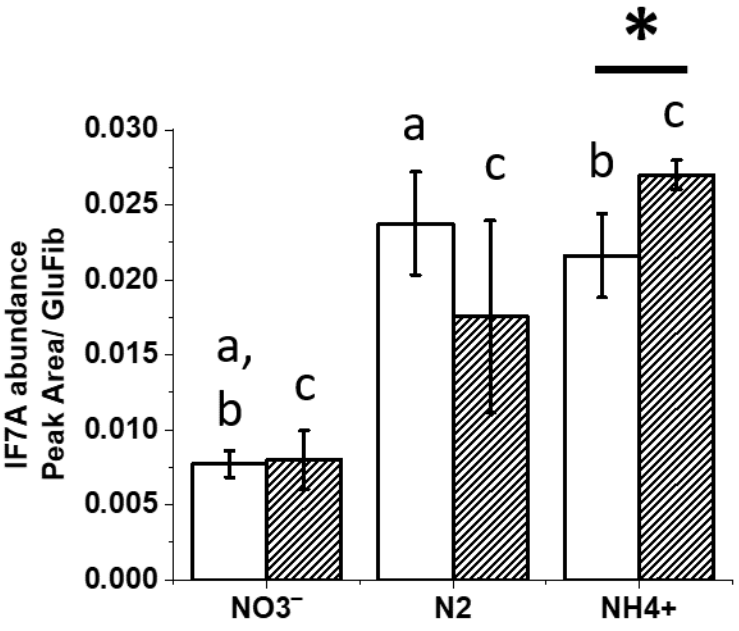


***Supplementary figure S3. Abundance of IF7A for 7120 WT and Δamt strains, after 96 h in the growth conditions of figure 2A.*** *Data are indicated as average ± SD of 6 biological replicates. Statistically significant differences between WT (white bars) and Δamt (striped bars) are indicated with an asterisk, whilst the same alphabet letter indicates statistically significant differences for the same strain in different growth conditions (one-way ANOVA, p-value < 0.05).*

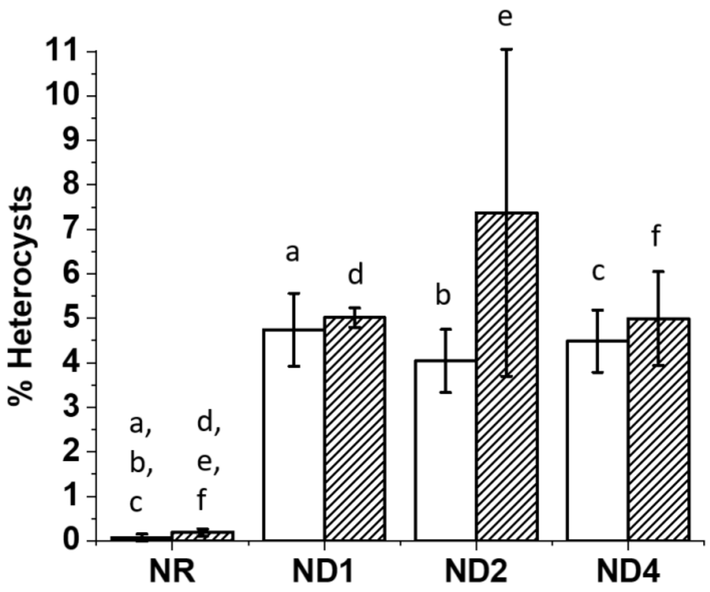


**Supplementary figure S4. Percentage of heterocysts in the filaments of both WT and *Δamt strains at different time points from N replete (NR) to N deplete conditions (ND)*.** *Time points are those indicated in Fig. 4A and correspond to NR (Nitrogen Replete), ND1, ND2 and ND4, respectively 1, 2 and 4 days after Nitrogen Deprivation. Data are expressed as percentage of heterocysts in the filament and are indicated as average ± SD of 3 biological replicates. For each biological replicate data come from the analysis of 10 microscope fields of view. See Supplementary Materials and Methods and supplementary figure S1 for information on the counting protocol. Statistically significant differences between WT (white bars) and Δamt (striped bars) are indicated with an asterisk, whilst the same alphabet letter indicates statistically significant differences for the same strain in different growth conditions (one-way ANOVA, p-value < 0.05).*

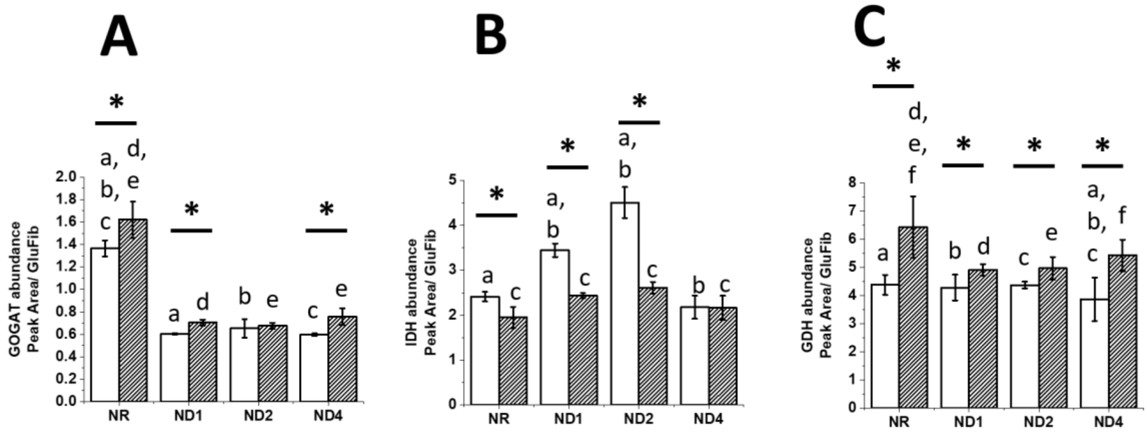


***Supplementary Figure S5. Abundance of GOGAT (A), IDH (B) and GDH (C) at different time points from N replete (NR) to N deplete conditions (ND) in both WT and Δamt strains.*** *Data are indicated as average ± SD of 6 biological replicates. Statistically significant differences between WT (white bars) and Δamt (striped bars) are indicated with an asterisk, whilst the same alphabet letter indicates statistically significant differences for the same strain in different growth conditions (one-way ANOVA, p-value < 0.05).*

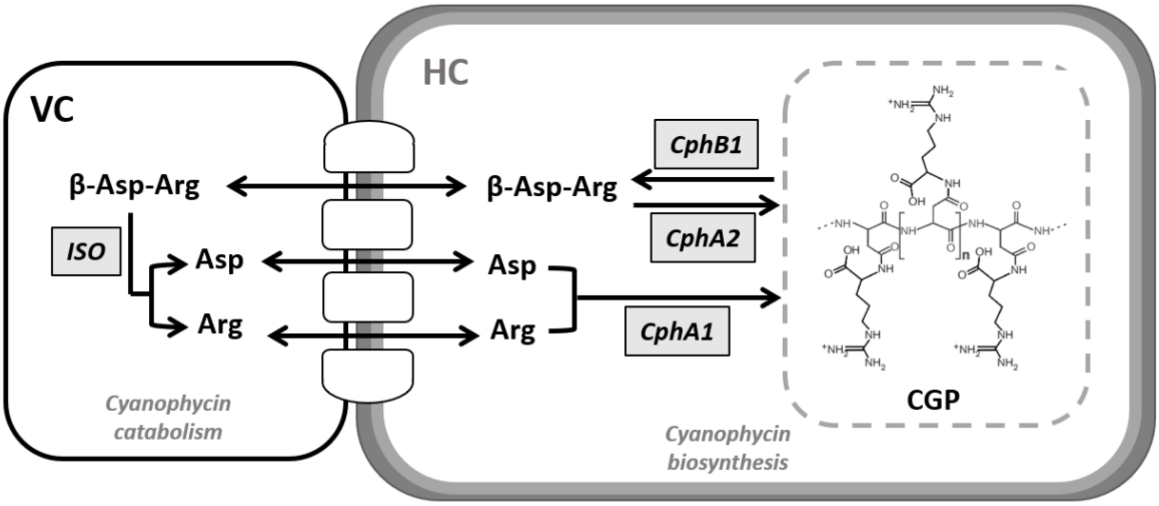


***Supplementary Figure S6. Schematic overview of cyanophycin granule polypeptide (CPG) metabolism in 7120, in diazotrophic conditions.*** *In 7120, CPG forms at the polar neck region of heterocysts (i.e. at their contact sites with the adjacent vegetative cells) and it is expected to regulate the transfer of fixed N from heterocysts (HC) to vegetative cells (VC). CPG is a polymer of the two amino acids aspartate (Asp) and arginine (Arg), forming a poly-L-aspartic acid backbone with each carboxyl group linked via isopeptide-bonds to an arginine residue (multi-L-arginyl-poly-L-aspartic acid). CPG biosynthesis: CPG is synthesised by a single enzymatic step catalysed by cyanophycin synthetase (i.e. CphA1), in a two-step reaction from L-aspartate and L-arginine, with one molecule of ATP involved in each step. CPG catabolism: CPG is first hydrolysed by cyanophycinase (i.e. CphB1), releasing the di-peptide β-aspartyl-arginine, which is consequently cleaved by isoaspartyl dipeptidase (i.e. ISO), generating the free amino acids Asp and Arg for further metabolic use. In 7120, as in many other N-fixing cyanobacteria, another cyanophycin synthetase (i.e. CphA2) is present, which uses the di-peptide β-aspartyl-arginine as substrate, directly recycling it into CPG. This scheme was adapted from (Forchhammer & Watzer, 2016).*

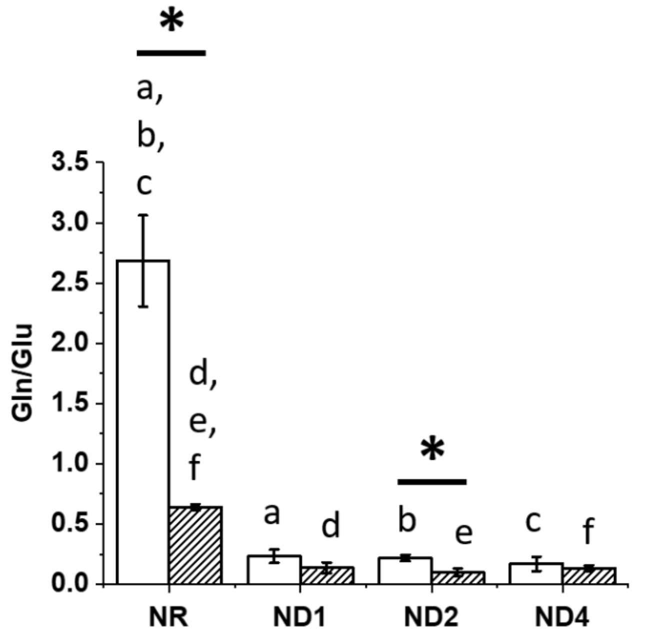


***Supplementary figure S7. Gln/Glu ratio in both WT (white bars) and Δamt strains (striped bars), grown in the experimental conditions of figure 4A.*** *Data are expressed as the ratio between Gln and Glu content in both strains as reported in Fig. 10A and 10B. Data are indicated as average ± SD of 6 biological replicates. Statistically significant differences between WT (white bars) and Δamt (striped bars) are indicated with an asterisk, whilst the same alphabet letter indicates statistically significant differences for the same strain in different growth conditions (one-way ANOVA, p-value < 0.05).*

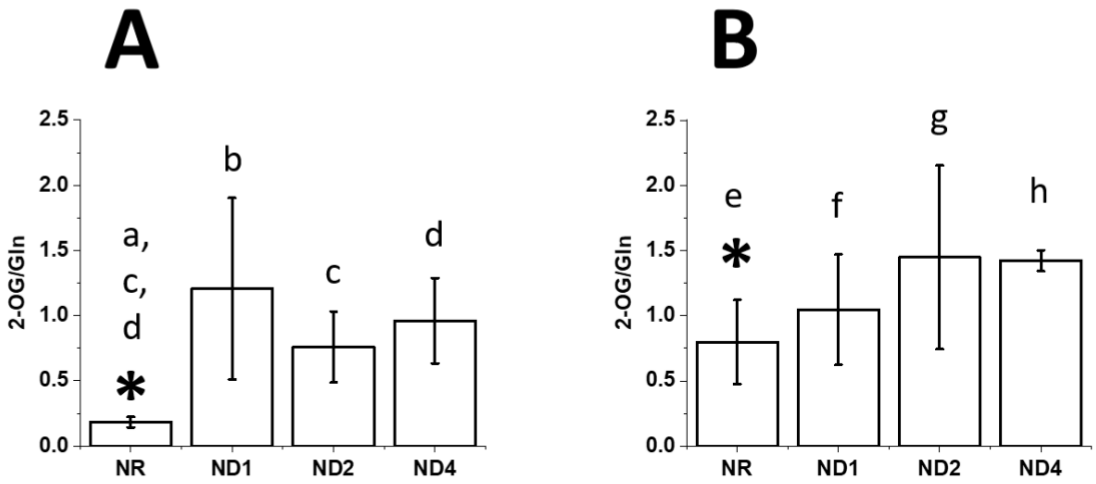


***Supplementary figure S8. 2-OG/Gln ratio in both WT (A) and Δamt strains (B), grown in the experimental conditions of figure 4A.*** *Data are expressed as the ratio between Gln and 2-OG content in both strains as reported in Fig. 10. Data are indicated as average ± SD of 6 biological replicates. Statistically significant differences between WT (A) and Δamt (B) are indicated with an asterisk, whilst the same alphabet letter indicates statistically significant differences for the same strain in different growth conditions (one-way ANOVA, p-value < 0.05).*
